# Oncogenic mutations convert MET from a pro-apoptotic tumor suppressor to an oncogenic driver

**DOI:** 10.1101/2025.11.10.687614

**Authors:** Rémi Tellier, Marie Fernandes, Audrey Vinchent, Sonia Paget, Agathe Laratte, Céline Vuillier, Elisabeth Werkmeister, Céline Villenet, Jean-Pascal Meneboo, Anne Chotteau-Lelièvre, Eric Wasielewski, Zoulika Kherrouche, Marie-José Truong, Clotilde Descarpentries, Luca Grumolato, Martin Figeac, Alexis B Cortot, Laurent Poulain, Andréa Paradisi, David Tulasne

## Abstract

Dependence receptors can exert both oncogenic and tumor-suppressive activities. In cancers, downregulation of dependence receptors or overexpression of their ligands are well-established mechanisms that drive tumor progression. However, direct genetic alterations abolishing the pro-apoptotic function of dependence receptors have not been documented so far. MET, a receptor tyrosine kinase classically viewed as an oncogene, has also been proposed to act as a dependence receptor through its caspase-mediated cleavage, but whether this property impacts tumorigenesis remained unknown. In ∼3% of lung adenocarcinomas, MET mutations leading to exon 14 skipping (METex14Del) remove both the caspase site and the adjacent CBL-binding motif, thereby preventing generation of the pro-apoptotic p40MET fragment. METex14Del promotes sustained signaling, enhanced invasion, apoptosis resistance, and tumor growth in HGF-humanized mice. Genome editing revealed that combined —but not individual— mutations of the caspase and CBL sites phenocopy METex14Del. Moreover, inducible re-expression of p40MET in METex14Del-expressing cells restored apoptosis and suppressed tumor formation. Altogether, our findings identify MET exon 14 skipping as the first oncogenic mutation that drives tumorigenesis by abolishing the tumor-suppressive pro-apoptotic function of a dependence receptor, thereby redefining the oncogenic potential of MET.

## INTRODUCTION

MET is a receptor tyrosine kinase (RTK) predominantly expressed in epithelial cells and activated by its high-affinity ligand, the hepatocyte growth factor (HGF). In mammals, the HGF/MET pair is essential for embryonic development^1–5^ and adult tissue regeneration^6–9^. MET activation involves homo-dimerization upon HGF binding to its extracellular domain, leading to intracellular kinase activation, downstream signaling, and cellular responses including proliferation, motility, and survival^10^.

The MET juxtamembrane domain contains negative regulatory sites, including a CBL (Casitas B-lineage Lymphoma) binding site (D_1002_Y_1003_R), whose tyrosine 1003, phosphorylated in response to HGF, plays a key role^11^. CBL is an E3 ubiquitin ligase involved in MET ubiquitination and degradation. Mutation of MET at Y1003 promotes motility in epithelial cells, focus formation in fibroblasts, and tumor growth in mice^11–13^. The CBL site partially overlaps with a caspase cleavage site (ESVD_1002_) responsible for MET cleavage during apoptosis. After a first caspase cleavage closer to the C-terminus, this cleavage produces a 40-kDa cytoplasmic C-terminal fragment (p40MET) which amplifies apoptosis by promoting mitochondrial permeabilization^14–19^.

Because MET induces survival upon ligand binding but apoptosis,-via caspase cleavage, in the absence of ligand, it has been classified as a dependence receptor (DR). As a consequence: knock-in mice expressing MET mutated at the caspase site showed resistance to apoptosis in epithelial organs such as the liver^16^. Many receptors are classified as DRs, including both RTKs, such as TrkC and c-KIT and non-RTKs, such as DCC and UNC5H^20–25^. DRs share two features: survival signaling in the presence of ligand and apoptosis in its absence. In the latter case, apoptosis is often due to a caspase-mediated cleavage converting the receptor to a pro-apoptotic factor.

Dependence receptors have been proposed to act as tumor suppressors^26^. Their expression, like that of other tumor suppressors, may be silenced in tumors, and this prevents apoptosis, as shown for the TrkC receptor^22^. Another mechanism through which DR pro-apoptotic function can be suppressed is sustained ligand stimulation, which inhibits caspase cleavage. For example, the TrkC ligand NT-3 promotes tumorigenesis by preventing TrkC-mediated apoptosis^27^. Conversely, use of a dominant-negative extracellular domain to block SCF binding to c-KIT promotes its cleavage and prevents tumor growth in mouse models^23^. Comparable inhibition of tumor growth has been seen after interfering with netrin binding to DCC and UNC5H^28^.

In all these examples, both activation of survival/proliferation/invasion pathways and inhibition of apoptosis are liable to account for the ligand’s pro-tumorigenic effects. However, as no direct molecular alterations specifically preventing DR cleavage have been clearly demonstrated in cancer, the role of impaired DR-mediated apoptosis in tumorigenesis remains insufficiently substantiated.

In non-small cell lung cancer (NSCLC), approximately three-quarters of patients harbor tumorigenesis-driving genomic alterations. Many of these are targetable by novel targeted therapies that improve survival and quality of life. Several alterations concern RTK-encoding genes and lead to direct kinase activation^29^. For the past 10 years, lung tumor profiling has revealed a group of MET-related alterations in about 3% of cases^30,31^. These tumors display alterations (called METex14Del) of splice sites of *MET* exon 14, causing in-frame exon 14 skipping and deletion of the receptor’s regulatory juxtamembrane domain. NSCLC patients with METex14Del show objective responses to MET-targeting TKIs, although response rates are lower (∼50%) than for other targetable alterations^32–35^.

Unlike activating mutations, MET exon 14 skipping removes a regulatory domain containing the CBL binding site and the caspase cleavage site, rather than directly activating the kinase. This makes it harder to elucidate the mechanisms of MET-induced activation and transformation in NSCLC^36^.

We previously demonstrated that METex14Del retains a strict requirement for its ligand HGF, not only to trigger cellular responses *in vitro* but also to sustain tumorigenic growth *in vivo*. This observation places METex14Del within the conceptual framework of dependence receptors, whose oncogenic activity remains contingent on ligand availability; yet, its potential pro-apoptotic function in the absence of ligand has remained unexplored in a cancer context^37^.

Here we demonstrate that MET exon 14 skipping prevents receptor cleavage by caspases and confers resistance to apoptosis. By using genome editing to inactivate individual juxtamembrane sites, we show that the caspase site specifically mediates apoptosis resistance. In HGF-humanized mice, we find this resistance to apoptosis to be required for MET-induced tumor growth. Yet, inducible re-expression of p40MET in METex14Del-expressing cells restored apoptosis and delayed tumor formation. Taking all this together, we evidence for the first time that the pro-apoptotic property of a dependence receptor can be overcome by cancer-associated mutations and that this mechanism is required for tumor growth.

## RESULTS

### MET exon 14 skipping induces sustained signaling, cell motility, and resistance to apoptosis

To investigate the impact of MET exon 14 skipping on downstream signaling activation and biological responses, we performed CRISPR-Cas9 genome editing on cells of the non-transformed lung epithelial cell line 16HBE14o-(16HBE). 16HBE cells express physiological levels of MET without overexpression or gene amplification, thus providing a model relevant to the context of METex14Del patients, who similarly do not exhibit MET overexpression or amplification. Exon 14 skipping was achieved by targeting the exon 14 donor splice site. Two METex14Del clones were analyzed. A control 16HBE cell line was generated by CRISPR-mediated homologous recombination with a single-stranded oligodeoxynucleotide (ssODN) containing silent mutations. MET protein expression levels and membrane localization were assessed by flow cytometry and showed comparable expression between cell lines expressing MET WT and METex14Del (Figure S1).

In MET WT 16HBE cells, HGF stimulation induced low levels of MET phosphorylation (on residues Y1234/1235 and Y1003), and also AKT and ERK phosphorylation, all of which decreased rapidly by 3 hours post-stimulation (Figures 1A-1D). In contrast, both METex14Del clones displayed higher MET, AKT, and ERK phosphorylation from 30 min post-stimulation to up to 8 hours. As expected, the METex14Del clones were not phosphorylated at residue Y1003 in the juxtamembrane domain. Total MET levels decreased after HGF stimulation in both MET WT and METex14Del clones (Figures 1A-1D). These results were confirmed in the lung adenocarcinoma cell line A549 edited to express the METex14Del receptor: MET WT A549 cells exhibited low MET, AKT, and ERK phosphorylation, decreasing by 3 hours post-stimulation, whereas METex14Del A549 cells displayed sustained phosphorylation of MET (at Y1234/1235 but not Y1003), AKT, and ERK from 30 minutes to 8 hours post-stimulation (Figure S2A). In a wound-healing assay, only HGF stimulation markedly enhanced the migration of 16HBE clones expressing METex14Del, as compared to their WT counterparts (Figure 1E). In a wound-healing assay on Matrigel, only METex14Del clones displayed invasive behavior in response to HGF (Figure 1F). In a scattering assay, only METex14Del clones displayed scattering upon HGF stimulation (Figure 1G).

**Figure 1.**
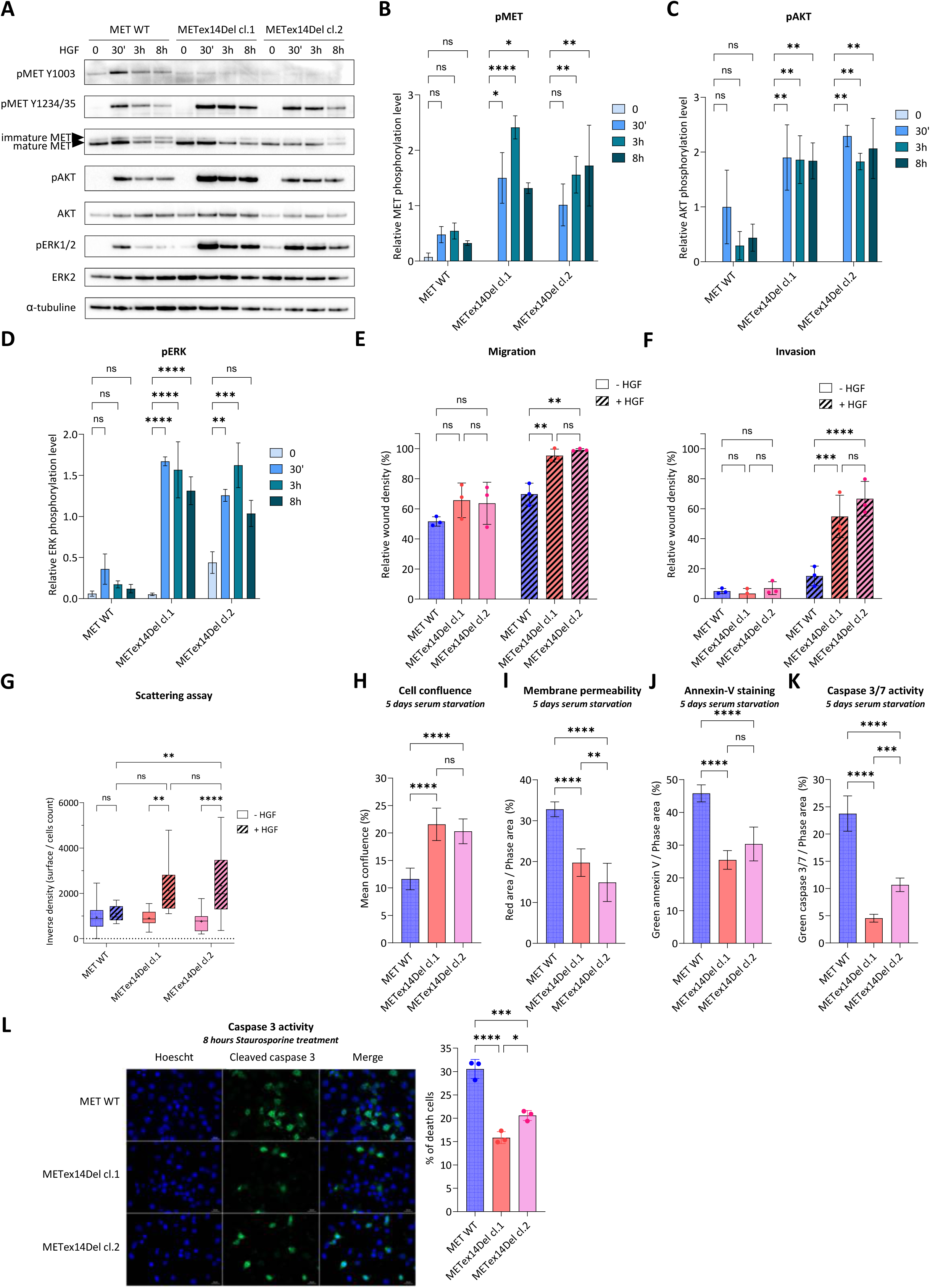
METex14Del induces sustained signaling, cell motility, and apoptosis resistance. (A) Immunoblot analysis of MET-induced downstream signaling pathways in edited 16HBE cells after 30 ng/mL HGF treatment for the indicated time. Immunoblotting is representative of three biological replicates. (B-D) Quantitative analysis of phosphorylated MET (p-MET), AKT (p-AKT), and ERK (p-ERK) normalized to their respective total protein levels from western blot experiments. Data are presented as means ± SEM from three independent experiments. Statistical significance was determined by two-way ANOVA followed by appropriate post hoc tests. ns (not significant), * (*p* < 0.05), ** (*p* < 0.01), *** (*p* < 0.001), **** (*p* < 0.0001). (E-F) Quantification of relative wound density after cell in the presence or absence of HGF in migration assays after 8 h post-HGF stimulation (E) and invasion assays with Matrigel 3 days post-HGF stimulation (F) following stimulation with 30 ng/mL HGF. Values represent mean ± SD from three independent experiments. Statistical significance was determined by two-way ANOVA followed by appropriate post hoc tests. ns (not significant), * (*p* < 0.05), ** (*p* < 0.01), *** (*p* < 0.001), **** (*p* < 0.0001). (G) Quantification of cell scattering based on cell density after 48 h stimulation with 30 ng/mL HGF. At least 15 cellular islets were analyzed. Data are presented as box-and-whiskers plots showing minimum to maximum values; in each box, the point represents the mean, and the horizontal line indicates the median. Statistical significance was determined using two-way ANOVA followed by appropriate post hoc tests. ns (not significant), ** (*p* < 0.01), **** (*p* < 0.0001). (H-K) Cell confluence (H), membrane permeability (I), Annexin V staining (J), and active caspase 3/7 staining (K) were measured five days after serum starvation. For (I–K), fluorescence signals were normalized to cell confluence. *n=6;* mean ± SD; representative of three independent experiments. Statistical significance was determined by one-way ANOVA. ns (not significant), ** (*p* < 0.01), *** (*p* < 0.001), **** (*p* < 0.0001). (L) Representative images showing nuclear staining with Hoechst (blue) and cleaved caspase 3 staining (green) after five hours of treatment with 1 µM staurosporine. Adjacent quantification shows the percentage of apoptotic cells, defined as cleaved caspase 3 positive cells with fragmented nuclei, from at least 200 counted cells per experiment. Data represent means ± SD from three independent experiments. Statistical significance was determined by one-way ANOVA. * (*p* < 0.05), *** (*p* < 0.001), **** (*p* < 0.0001).

To determine the transcriptomic program of 16HBE induced by ligand-stimulated METex14Del, we performed RNA sequencing following HGF stimulation. The two independent METex14Del clones exhibited a broad transcriptional profile in response to HGF, which supports the robustness of the phenotype induced by exon 14 skipping (Figure S3A). In contrast to MET WT cells, which displayed minimal transcriptomic changes upon HGF exposure, both METex14Del clones showed strong upregulation of gene sets associated with cell motility, extracellular matrix organization, angiogenesis, and signaling, as revealed by gene ontology analysis. This is in keeping with the biological phenotypes observed (Figures S3B-S3F).

To evaluate the impact of MET exon 14 skipping on apoptosis resistance, we performed a five-day serum starvation (at 0.5% FBS, unstimulated by HGF) experiment. This induced apoptosis in MET WT 16HBE cells, as evidenced by reduced confluence, increased Annexin V staining, membrane permeability, and caspase 3/7 activity. All these responses were reduced in cells expressing METex14Del (Figure 1H-1K). Similar resistance to serum-starvation-induced apoptosis was observed in METex14Del A549 cells (Figure S2B-S2D). In both 16HBE and A549 cells, METex14Del also conferred partial resistance to staurosporine-induced short-term apoptosis (Figures 1L and S4A-S4E).

Collectively, these findings reveal a dual METex14Del function: on the one hand and consistently with the observed transcriptional program, stimulation by HGF of METex14Del induces sustained downstream signaling promoting increased cell migration, invasion, and scattering. On the other hand, METex14Del confers intrinsic resistance to apoptosis, without ligand stimulation.

### Capmatinib does not prevent METex14Del-induced tumor growth from 16HBE cells

Given the enhanced signaling and motility induced by METex14Del upon HGF stimulation, we assessed how effectively the MET-selective tyrosine kinase inhibitor capmatinib might suppress these responses. Capmatinib abolished MET, AKT, and ERK phosphorylation in both MET WT and METex14Del 16HBE cells (Figure 2A). Results were similar with the A549 cell line (Figure S5A). Consistently, capmatinib blocked METex14Del-driven migration and invasion in response to HGF (Figures 2B and 2C). By contrast, it had no effect on METex14Del-mediated resistance to apoptosis, as attested by comparable levels of membrane permeability during serum starvation. The apoptosis resistance phenotype is thus independent of MET kinase activity (Figures 2D and S5B).

**Figure 2.**
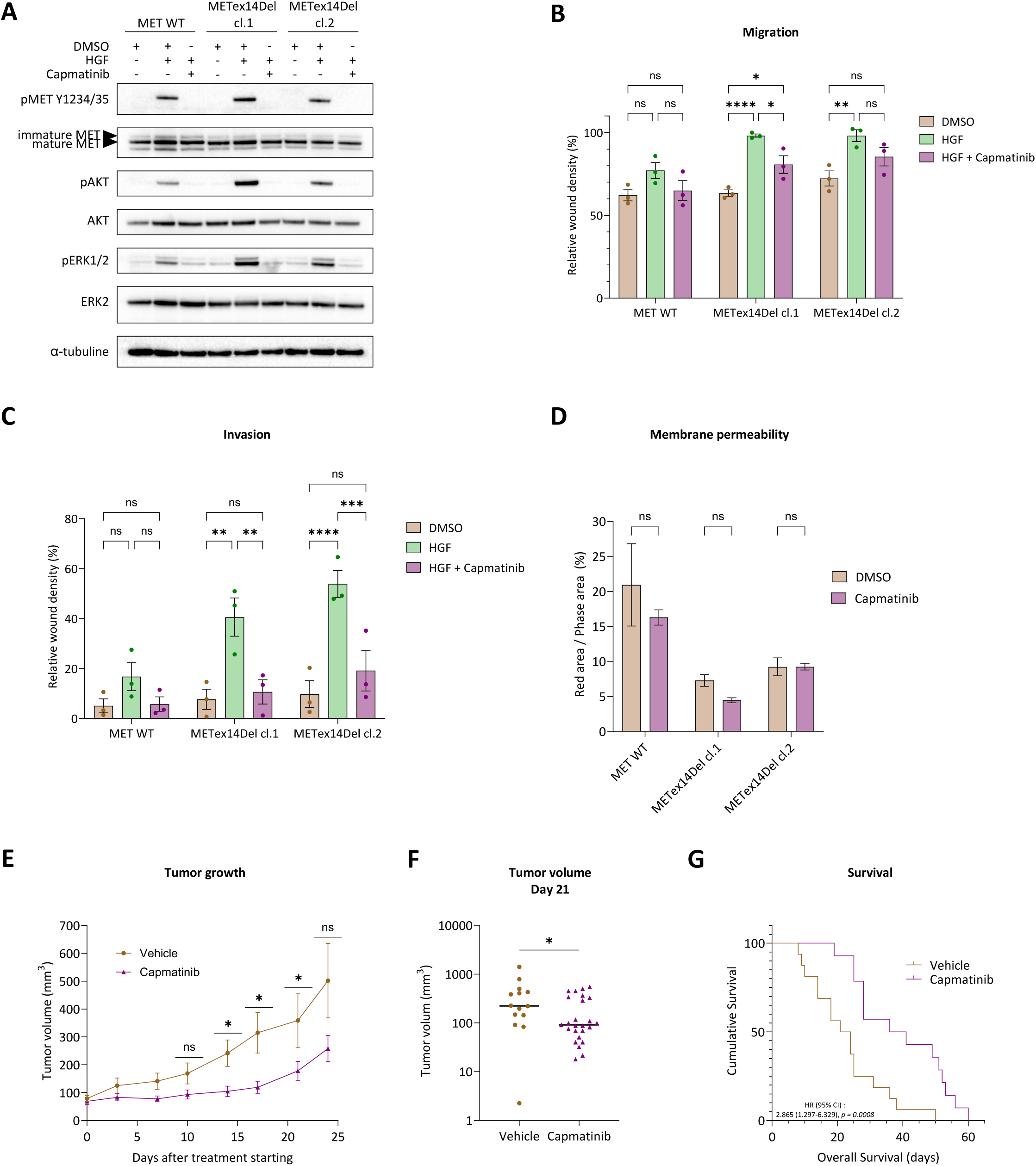
Capmatinib partially inhibits METex14Del-induced tumor growth in 16HBE cells. (A) Immunoblot analysis of MET-induced downstream signaling pathways in edited 16HBE cells treated with 30 ng/mL HGF for 30 min, with or without capmatinib pre-treatment. Cells were pre-treated with 1 µM capmatinib for 90 minutes prior to HGF stimulation. Immunoblots are representative of three independent biological experiments. (B-C) Quantification of relative wound density in migration assays (8 h post-HGF stimulation) and invasion assays (in Matrigel, 3 days post-HGF stimulation) following stimulation with 30 ng/mL HGF, with or without 1 µM capmatinib pre-treatment for 90 minutes. Values represent means ± SD from three independent experiments. Statistical significance was determined by two-way ANOVA followed by appropriate post hoc tests. ns (not significant), * (p < 0.05), ** (p < 0.01), *** (p < 0.001), **** (p < 0.0001). (D) Membrane permeability was measured five days after serum starvation with or without 1 µM capmatinib. The fluorescence signal was normalized to cell confluence. *n=5;* mean ± SD; representative of two independent experiments. Statistical significance was determined by one-way ANOVA. ns (not significant). (E-G) Tumor growth in and survival of NSG-huHGF mice injected subcutaneously with genome-edited cells. Daily treatment with vehicle or capmatinib (10 mg/kg/day) was initiated when at least one tumor per mouse reached approximately 100 mm³. (E) Tumor growth curves from the start of treatment. Data represent mean tumor volumes ± SEM. (F) Tumor volume at day 21; each point represents an individual tumor. (G) Kaplan–Meier curve showing overall survival. Statistical significance was assessed with unpaired two-tailed *t*-tests for tumor volume comparisons (E, F) and the log-rank (Mantel–Cox) test for survival (G). The hazard ratio (HR) is indicated. ns (not significant), * (p < 0.05).

Next, METex14Del 16HBE cells were xenografted into immunodeficient mice expressing human HGF (NSG-huHGF) (which in contrast to murine HGF, is able to activate the human MET expressed in xenografted cells). As METex14Del activation depends on its ligand (consistently with the behavior of dependence receptors), the use of human-HGF-expressing mice was required. In these mice, daily capmatinib treatment partially inhibited tumor growth: after three weeks of treatment, tumor volumes were reduced as compared to the vehicle-only group, and this led to increased overall survival (Figures 2E–2G). In SCID mice, in contrast, capmatinib treatment drastically inhibited tumor growth from xenografted Hs746T gastric carcinoma cells harboring both the METex14Del mutation and MET gene amplification (leading to ligand-independent receptor activation, as confirmed by a signaling study (Figure S6A)). This demonstrates the efficacy of a MET TKI in a situation where oncogene addiction is dependent on MET kinase activity (Figure S6B). Capmatinib inhibited MET, AKT, and ERK phosphorylation *in vitro* in these cells (Figure S6B).

These results demonstrate that, while MET TKI effectively blocks METex14Del-mediated signaling and motility *in vitro*, it fails to completely inhibit tumor growth. This suggests that kinase-independent mechanisms may support tumorigenesis in this context.

### Landscape of MET exon 14 mutations in a large NSCLC cohort

To comprehensively characterize the spectrum of MET exon 14 alterations in non-small cell lung cancer (NSCLC), we analyzed formalin-fixed, paraffin-embedded (FFPE) tumor samples from 6,990 patients referred for routine molecular screening between May 2017 and December 2024. Targeted next-generation sequencing was performed with the optimized CLAP panel, allowing full coverage of MET exon 14, its flanking intronic regions, and other clinically relevant alterations in lung cancer^37^. Within the total cohort, the most frequent oncogenic drivers were mutations in *KRAS* (36.2%), *EGFR* (12.3%), *BRAF* (5.4%), and *ERBB2* (<1%). MET exon 14 splice-site alterations —predicted to induce exon skipping— were identified in 166 cases (2.4%), representing 93.2% of all MET alterations (178/6990). Beyond these canonical splice variants, 12 additional mutations or in-frame deletions were detected within the coding region of exon 14. These rare events led to predicted amino-acid changes including p.Tyr971Cys, p.Val975-Ser985del, p.His979Arg, p.Ser985Thr, p.Leu983Phe, p.Glu999Lys, p.Asp1002Tyr, p.Asp1002-Thr1006del, p.Tyr1003Asn and p.Tyr1003del. Notably, 7 of these 12 variants (3.4% of the MET exon 14 mutations) were predicted to affect known regulatory sites (Ser985, Tyr1003 and Asp1002). These findings broaden the mutational spectrum of MET exon 14 beyond canonical splice-site events. Although exon 14 skipping represents the dominant alteration in NSCLC, we also identify rare coding mutations affecting notably the CBL-binding and caspase-cleavage sites. Their occurrence, despite low frequency, suggests that loss of these regulatory checkpoints may be functionally important. This raises the possibility that disruption of either or both sites contributes to MET-driven tumorigenesis and warrants functional investigation of their individual and combined loss (Table 1).

**Table 1.** Molecular Landscape of MET Exon 14 Alterations in 6990 NSCLC patients.

| Molecular alteration localization<br>from 5' to 3' end | Recurrence (n=178) | Expected consequences |
| --- | --- | --- |
| Exon 14 splicing site | 166/178 | Exon 14 skipping |
| Exon 14 c.3049G>A | 1/178 | p.Glu999Lys |
| Exon 14 c.3059_3070del | 1/178 | p.Asp1002_Thr1006delinsAla |
| Exon 14 c.3061_3063del | 1/178 | p.Tyr1003del |
| Exon 14 c.3061T>A | 2/178 | p.Tyr1003Asn |
| Exon 14 c.3001C>T | 2/178 | p.Leu983Phe |
| Exon 14 c.3058G>T | 1/178 | p.Asp1002Tyr |
| Exon 14 c.3008G>C | 1/178 | p.Ser985Thr |
| Exon 14 c.2990A>G | 1/178 | p.His979Arg |
| Exon 14 c.2978_3010del | 1/178 | p.Val975_Ser985del |
| Exon 14 c.2966A>G | 1/178 | p.Tyr971Cys |

### Loss of the CBL binding and caspase cleavage sites does not recapitulate the METex14Del-driven signaling and cell responses

To determine the respective contributions of the CBL binding site and the caspase cleavage site to METex14Del-driven phenotypes, we performed genome editing to generate 16HBE cell lines harboring the MET Y1003F mutation, the MET V1001A mutation, or both (Figure 3A). As the D1002 residue is shared by both sites^18^, the V1001 residue was targeted in order to selectively disrupt the caspase site. All edited cell lines displayed the expected mutation(s), as confirmed by sequencing of MET exon 14 (Figure S7). MET protein levels and membrane localization were unaffected as assessed by flow cytometry (Figure S1).

**Figure 3.**
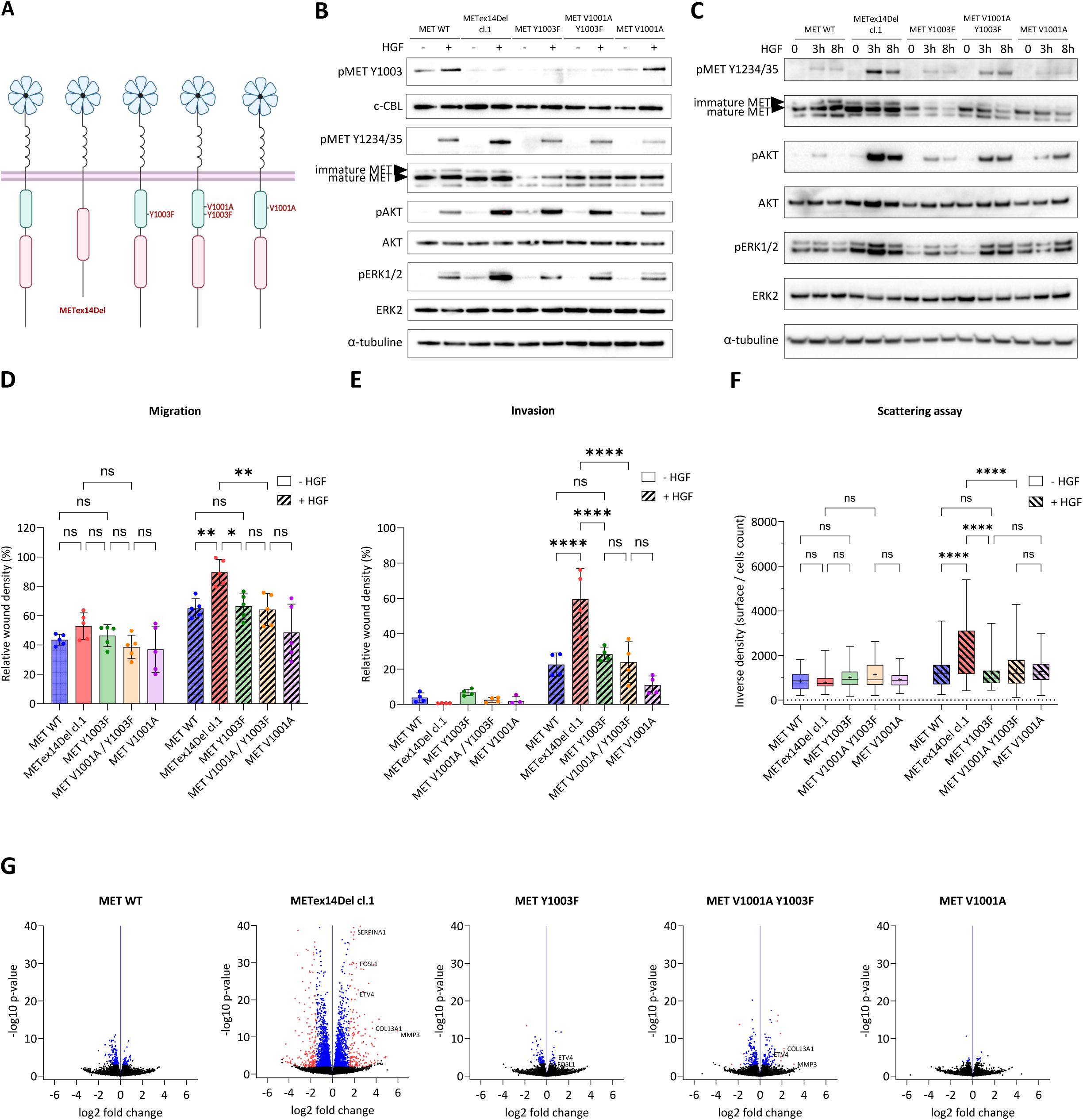
Loss of the CBL and caspase sites does not recapitulate the sustained signaling and biological responses of METex14Del. (A) Schematic representation of the five 16HBE clones generated by CRISPR-Cas9 genome editing. Mutations identified in each clone are highlighted in red. (B-C) Immunoblot analysis of MET-induced downstream signalling pathways in edited 16HBE cells after 30 ng/mL HGF treatment for the indicated time. Immunoblotting is representative of three biological replicates. (D-E) Quantification of relative wound density in migration assays (D) (without Matrigel, 8 h post-HGF stimulation) and invasion assays (E) (with Matrigel, 3 days post-HGF stimulation) following stimulation with 30 ng/mL HGF. Values represent mean ± SD from four or fiive independent experiments. Statistical significance was determined by two-way ANOVA followed by appropriate post hoc tests. ns (not significant), * (*p* < 0.05), ** (*p* < 0.01), *** (*p* < 0.001). (F) Quantification of cell scattering based on the inverse of cell density after 48 h stimulation with 30 ng/mL HGF. At least 40 cellular islets were analyzed. Data are presented as box-and-whiskers plots showing minimum to maximum values; in each box the point represents the mean and the horizontal line indicates the median. Statistical significance was determined by two-way ANOVA followed by appropriate post hoc tests. ns (not significant), **** (*p* < 0.0001). (F) Volcano plots of RNA sequencing data from genome-edited 16HBE clones showing differential gene expression relative to control samples. Each point represents a gene, plotted as –log₁₀(*p*-value) versus log₂ fold change. Genes with adjusted *p*-value (BH) < 0.05 are shown in blue. Genes with adjusted *p*-value (BH) < 0.05 and absolute fold change greater than 1.5 value are highlighted in red.

Upon HGF stimulation, phosphorylation at Y1003 was detected in MET WT and MET V1001A cells, but not in MET Y1003F, MET V1001A Y1003F, or METex14Del cells. This confirms the integrity of this phosphorylation site in MET V1001A cells. None but the METex14Del cells showed, upon HGF stimulation, the above-mentioned increased and sustained MET, AKT, and ERK phosphorylation (observed in Y1003F-mutated cell lines at 30 min but maintained only in METex14Del cells at 8 hours) and increased migration, invasion, and scattering (Figure 3B-3F).

Furthermore, only METex14Del cells exhibited a robust transcriptional response, characterized by widespread gene up and down-regulation. In contrast, consistently with their limited biological responses to HGF, the MET Y1003F, MET V1001A, and double mutant displayed transcriptomic profiles closely related to that of MET WT, with minimal gene expression changes following HGF exposure (Figure 3G). Consistently with the gene ontology term enrichments observed for METex14Del clones (Figure S3B), several key genes involved in extracellular matrix remodeling, migration and signaling — including *MMP3*, *SERPINB5*, *SERPINB8*, and *FOSL1*— were specifically upregulated in response to HGF in METex14Del cells, but not in any other mutant cell lines (Figures S8A–S8D). Taken together, our results indicate that disruption of either the CBL binding site, the caspase cleavage site, or both does not recapitulate the phenotype (sustained signaling, motility, transcriptional program) induced by HGF in METex14Del-expressing cells.

### Loss of the caspase cleavage site confers, via impaired p40MET generation, resistance to serum-starvation-induced apoptosis

Given the observed resistance of METex14Del cells to apoptosis, we investigated whether this phenotype results from loss of the caspase cleavage site. First, an *in vitro* cleavage assay with purified caspase 3 was applied to lysates of the WT and mutant cells. After treatment with caspase 3, we probed MET-mutated proteins with two antibodies: MET-KC, recognizing the kinase-proximal C-terminal region and detecting MET even after caspase-mediated C-terminal cleavage, and MET-EC, recognizing the extreme C-terminal tail, which does not detect the cleaved receptor. MET-KC antibody revealed that the pro-apoptotic fragment p40MET was detected in the MET WT lysate and at a lower level in the MET Y1003F lysate, but not in the lysates of cells mutated at the juxtamembrane caspase site (V1001A Y1003F and V1001A) (Figure 4A). As expected, caspase 3 cleaved MET c-terminally with respect to p40MET, as revealed with the MET-EC antibody. PARP1/2, used as a control, showed the expected cleavage in the presence of caspase 3. The same assay applied to A549 cell lysates gave similar results (Figure S9). The p40MET fragment was also generated during staurosporine-triggered apoptosis in MET WT and Y1003F cells, but not in the caspase-site mutants (Figure S10).

**Figure 4.**
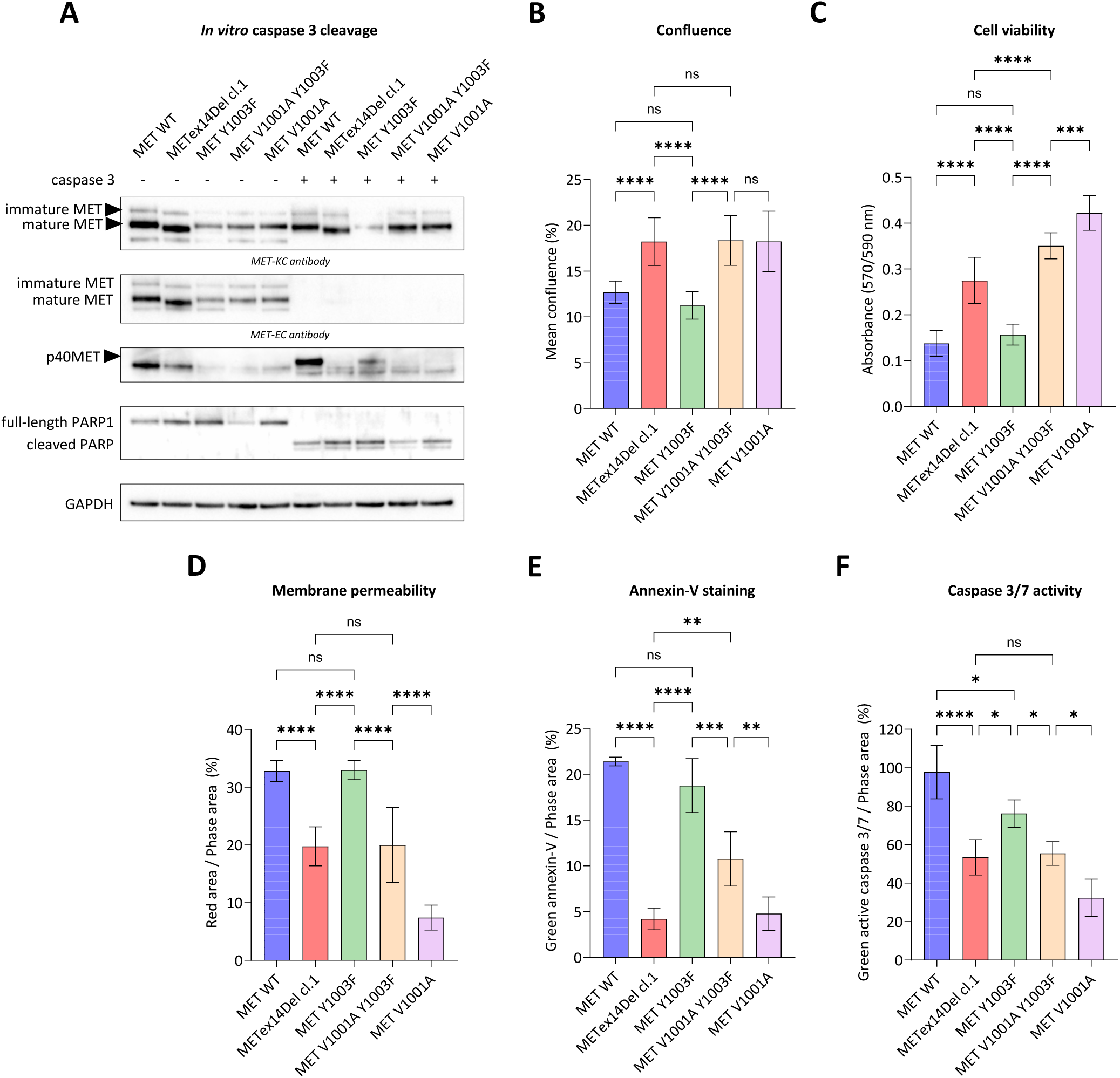
Loss of the caspase site confers resistance to serum starvation-induced apoptosis via impaired p40MET generation. (A) Immunoblot analysis showing generation of the p40MET fragment following *in vitro* cleavage of MET by recombinant caspase 3 in 16HBE cell lysates. MET-mutated proteins level was analyzed using two anti-MET antibodies: MET-KC, recognizing the kinase-proximal C-terminal region and detecting MET even after caspase-mediated C-terminal cleavage, and MET-EC, recognizing the extreme C-terminal tail, which does not detect the cleaved receptor. Immunoblots are representative of three independent biological experiments. (B-F) Cell confluence (B), viability (C), membrane permeability (D), Annexin V staining (E), and active caspase 3/7 staining (F) were measured five days after serum starvation. For (C), viability was measured five days after serum starvation in an AlamarBlue assay. For (D–F), fluorescence signals were normalized to cell confluence. *n=6;* mean ± SD; representative of three independent experiments. Statistical significance was determined by one-way ANOVA. ns (not significant), * (*p* < 0.05), ** (*p* < 0.01), *** (*p* < 0.001), **** (*p* < 0.0001).

To functionally assess the impact of caspase site loss, 16HBE cells were grown for five days under serum starvation. While MET Y1003F cells remained sensitive to serum starvation, both MET V1001A Y1003F and MET V1001A mutants phenocopied the resistance to apoptosis of METex14Del expressing cells, as evidenced by higher confluence and viability, reduced membrane permeability, lower Annexin V staining, and decreased caspase 3 activity (Figures 4B–4F).

To rule out the possibility that the observed differential resistance to apoptosis might have arisen from variations in apoptosis regulator expression, we assessed the levels of key apoptosis-related proteins across all cell lines under standard culture conditions. Expression of the BH3-only proteins (Bim, Bid, Puma, Noxa), of the pro-apoptotic effectors Bax and Bak, and of the anti-apoptotic proteins Bcl-x_L_ and Mcl-1 remained largely unchanged. It is worth noting that Bcl-2 was undetectable in all cell lines (Figures S11A-S11I).

Taken together, our results indicate that failure to generate p40MET due to disruption of the caspase cleavage site, is sufficient to confer resistance to serum-starvation-induced apoptosis. The fact that this resistance to apoptosis occurs in the absence of HGF stimulation suggests that it is independent of kinase activation. This highlights a kinase-independent survival function of METex14Del, associated with impaired caspase cleavage of the receptor.

### Combined loss of the CBL binding site and caspase cleavage site recapitulates METex14Del-driven tumorigenesis

To assess the tumorigenic potential of MET exon 14-associated mutations, the different genome-edited 16HBE cell lines were subcutaneously injected into NSG-huHGF mice. When either MET WT, MET Y1003F, or MET V1001A cells were injected at 14 sites, in each case, only one small tumor was observed 59 days post-injection. In contrast, METex14Del and MET V1001A Y1003F cells each gave rise to tumors at 13 out of 14 injections, with comparable mean tumor volumes (411.6 mm³ and 382.9 mm³, respectively) (Figures 5A–5C). Tumors derived from METex14Del and MET V1001A Y1003F cells expressed membrane-located human MET protein (Figure 5D). Proliferation within the tumor tissue was confirmed by PCNA staining (Figure 5E).

**Figure 5.**
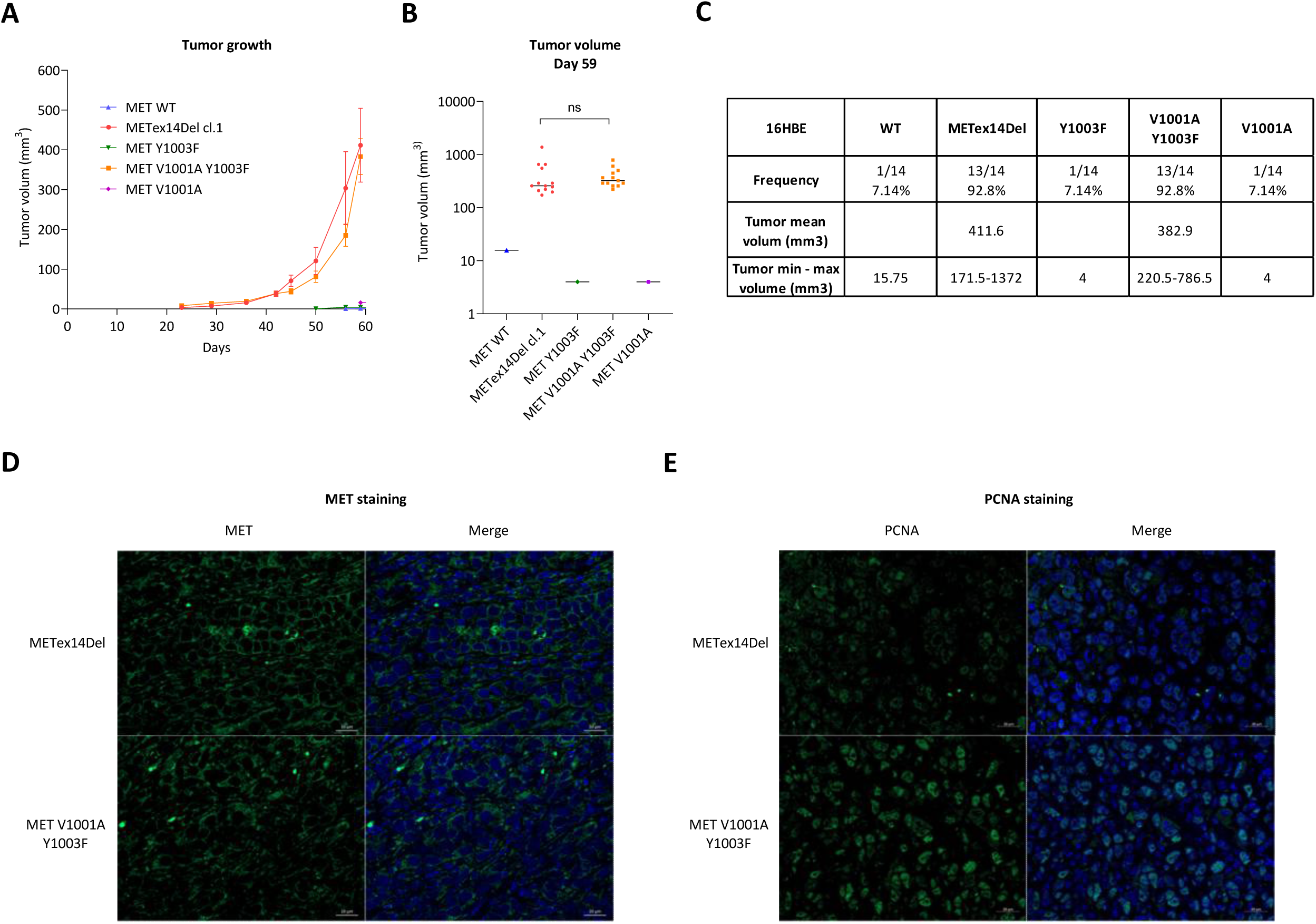
Combined loss of CBL binding and caspase cleavage sites recapitulates METex14Del-driven tumorigenesis. (A-C) Tumor growth in NSG-huHGF mice injected subcutaneously with genome-edited cells. (A) Tumor growth curves from the time of cell injection. Data represent mean tumor volume ± SEM. (B) Tumor volume at day 59; each point represents an individual tumor. (C) Frequency and mean tumor volume for each edited 16HBE clone after 59 days. Statistical significance was assessed with unpaired two-tailed *t*-tests for tumor volume comparisons. ns (not significant). (D) IHC for MET expression (green) in tumor sections, with Hoechst nuclear counterstaining (blue). (E) IHC for the proliferation marker PCNA (green), with Hoechst nuclear counterstaining (blue).

These results suggest that METex14Del-driven tumorigenesis relies on the loss of both the CBL binding site and the caspase cleavage site, conferring two complementary properties, including HGF-independent resistance to apoptosis.

### Induced p40MET expression restores sensitivity to apoptosis and inhibits METex14Del-driven tumorigenesis

Given the effect of caspase site loss on METex14Del-induced tumorigenesis, we investigated how pro-apoptotic p40MET fragment might affect tumorigenesis by engineering METex14Del 16HBE cells to express the fragment under the control of doxycycline (Figure 6A). Induced p40MET expression did not trigger apoptosis in cells cultured under standard serum conditions (as evidenced by the absence of PARP1/2 cleavage (Figure 6A)). Nor did it modify METex14Del-mediated HGF-dependent downstream signaling (Figure 6B) or cell migration and invasion (Figures 6C and 6D). In contrast, p40MET expression in METex14Del cells restored sensitivity to long-term apoptosis induced by serum starvation, an effect not observed in doxycycline-treated control cells (Figure 6E).

**Figure 6.**
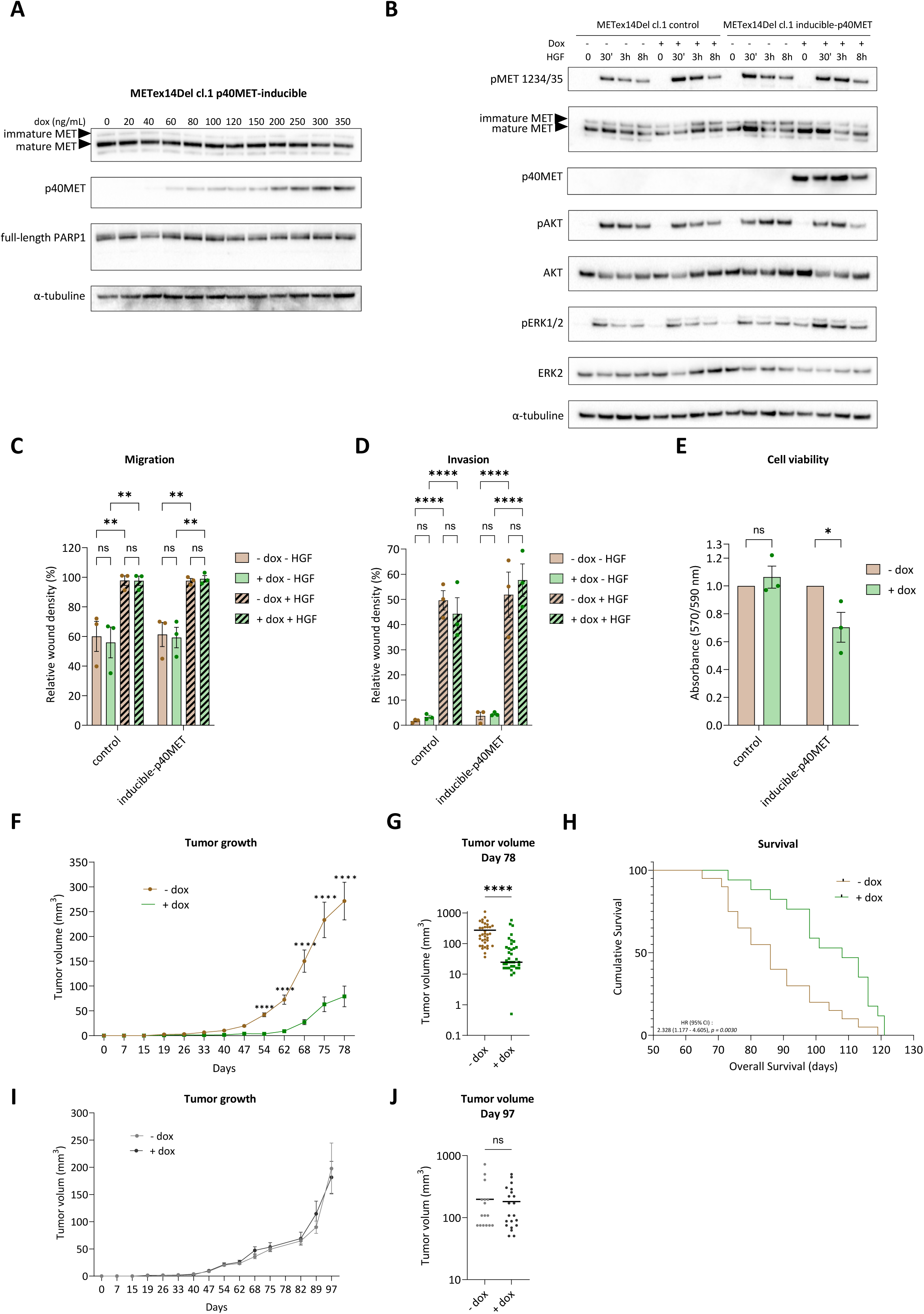
Induced p40MET expression restores sensitivity to apoptosis and inhibits METex14Del-driven tumorigenesis. (A) Immunoblot analysis of inducible p40MET fragment expression in METex14Del cl.1 16HBE cells after 16 h of doxycycline treatment at the indicated concentration. (B) Immunoblot analysis of MET-induced downstream signaling pathways after treatment with 30 ng/mL HGF for the indicated time with or without overnight doxycycline pre-treatment at 1 µg/mL. Immunoblotting is representative of two biological replicates. (C-D) Quantification of relative wound density in migration assays (C) (without Matrigel, 8 h post-HGF stimulation) and invasion assays (D) (with Matrigel, 3 days post-HGF stimulation) following stimulation with 30 ng/mL HGF, with or without overnight doxycycline pre-treatment at 1 µg/mL. Values represent means ± SD from three independent experiments. Statistical significance was determined by two-way ANOVA followed by appropriate post hoc tests. ns (not significant), ** (*p* < 0.01), *** (*p* < 0.001). (E) Cell viability was measured five days after serum starvation in an AlamarBlue assay. Values represent means ± SD from three independent experiments. Statistical significance was determined by two-way ANOVA followed by appropriate post hoc tests. ns (not significant), * (*p* < 0.05). (F-H) Tumor growth in NSG-huHGF mice injected subcutaneously with 16HBE METex14Del cells inducibly expressing p40MET. (F) Tumor growth curves from the time of cell injection. Data represent means tumor volumes ± SEM. (G) Tumor volume at day 78; each point represents an individual tumor. (H) Kaplan–Meier curve showing overall survival. Statistical significance was assessed with unpaired two-tailed *t*-tests for tumor volume comparisons (F, G) and with a log-rank (Mantel–Cox) test for survival (H). The hazard ratio (HR) is indicated. **** (*p* < 0.0001). (I–J) Tumor growth in mice subcutaneously injected with METex14Del 16HBE cells. Doxycycline (1 mg/mL) was provided in the drinking water from the time of cell injection. (I) Tumor growth curves from the day of injection. Data represent means tumor volumes ± SEM. (J) Tumor volumes at day 97; each point corresponds to an individual tumor. Statistical significance was assessed with unpaired two-tailed *t*-tests. ns, not significant.

To determine the consequence of this observation on tumorigenesis, p40MET inducible METex14Del cells were injected subcutaneously into NSG-huHGF mice. Doxycycline-induced p40MET expression markedly inhibit tumor growth, with mean tumor volume of 271 mm³ with vehicle versus 76 mm³ in doxycycline-treated mice (Figure 6F). Moreover, p40MET expression significantly extended overall survival (Figure 6H). Importantly, doxycycline treatment had no effect on tumor growth from non-transfected METex14Del control cells (Figures 6I and 6J). Together, these results demonstrate that restoring p40MET expression is sufficient to resensitize METex14Del cells to apoptosis and to slow down tumor growth.

## DISCUSSION

RTKs are predominantly known to be oncogenic, being frequently activated by gain-of-function mutations, gene amplification, or chromosomal rearrangements leading mainly to ligand-independent signaling driving tumorigenesis^38^. The concept of “oncogene addiction” has motivated the development of targeted therapies, particularly TKI therapy, for many cancers^39^. Yet, a growing body of evidence justifies a more nuanced view: some RTKs play a dual role, acting also as dependence receptors. These receptors promote cell survival when ligand-bound, but can induce apoptosis in its absence, thus functioning as conditional tumor suppressors^26^. TrkC provides an illustrative example of this duality: in colorectal cancer, loss of TrkC expression through promoter methylation, effectively prevents it from exerting its pro-apoptotic function^22^ whereas in cancers such as breast cancer or melanoma, overexpression of its ligand NT-3 inhibits TrkC-mediated apoptosis without any genetic alteration of the receptor itself^21^. Similarly, c-KIT requires binding to SCF to suppress apoptosis, and disruption of this ligand-receptor interaction enhances apoptosis and limits tumor growth^23^.

Importantly, while activating mutations conferring oncogenic gain-of-function are common and well-characterized in RTKs, direct mutations specifically abolishing their apoptosis-promoting, tumor-suppressive functions are undocumented. Instead, loss of such an apoptosis-checkpoint in cancer mostly results from decreased receptor expression or elevated ligand availability, and this suggests a regulatory balance rather than an on/off genetic switch. Thus, whereas classical tumor suppressors are often inactivated by mutations, RTK-mediated tumor suppression via a dependence receptor function is primarily bypassed through epigenetic silencing or ligand-mediated inhibition.

This duality raises an important question: while RTK-activating mutations are typically classified as oncogenic because of their ability to drive constitutive signaling and tumorigenesis, they may simultaneously abolish a latent tumor-suppressive, pro-apoptotic function. Yet, this loss is often overlooked —concealed within the dominant gain-of-function phenotype— because the amplified survival and proliferation signals mask any disruption of apoptotic signaling. Recognizing and dissecting this layered effect is critical, as it may reveal previously unappreciated vulnerabilities and open new avenues for targeting RTKs beyond their canonical oncogenic activity.

MET exemplifies this dual-function paradigm. While ligand-bound MET promotes proliferation, survival, and migration, ligand withdrawal triggers caspase-mediated cleavage at a juxtamembrane site, generating a pro-apoptotic intracellular fragment (p40MET) which amplifies mitochondrial apoptosis^17^. This cleavage has been validated *in vitro* and *in vivo*, with non-cleavable MET mutants conferring apoptosis resistance, notably in the liver^16^. Despite these features, the contribution of MET’s apoptosis-promoting function to tumor suppression has remained largely unexplored. We have addressed this question using the clinically relevant MET exon 14 skipping mutation (METex14Del), found in ∼3% of non-small cell lung cancers. This alteration removes the juxtamembrane domain, eliminating both the CBL-binding site and the caspase cleavage site, and is known to promote sustained MET signaling^30^. However, its potential impact on apoptosis regulation has not been previously examined.

Using human bronchial epithelial cells (16HBE), devoid of any other oncogenic alterations, we have established a physiologically relevant model for dissecting the specific contribution of METex14Del. This approach has enabled us to isolate the effects of this mutation in the absence of other, confounding molecular alterations. This is essential as dependence receptor signaling relies on a finely tuned balance between survival and apoptotic cues. In this model, we found METex14Del to remain strictly ligand-dependent for all canonical oncogenic outputs, including signaling, migration, invasion, and cell scattering. In contrast, the apoptotic resistance conferred by METex14Del occurred independently of HGF stimulation, consistently with the defining feature of dependence receptors. The robustness and relevance of this mechanism is supported by the fact that ligand-independent apoptotic protection was also observed in the lung cancer cell line A549 displaying MET exon 14 skipping. Furthermore, the fact that METex14Del drives tumorigenesis only in the presence of its ligand prompted us to utilize HGF-humanized mice to accurately model this dependency *in vivo*^37^.

A unique feature of MET is that its CBL-binding motif (D^1002^Y^1003^R) and its caspase cleavage site (D1002) are juxtaposed within the juxtamembrane domain, forming a regulatory hub where phosphorylation and cleavage are mutually exclusive: phosphorylation of Y1003 prevents caspase access, while cleavage detaches the kinase domain from the receptor, thereby rendering it inactive^18^. This structural arrangement strongly supports a ligand-gated molecular switch consistent with the dependence receptor model. To decode the specific contributions of each motif, we used precise genome editing to selectively disrupt the caspase site or the CBL-binding site independently.

Our results demonstrate that loss of the caspase cleavage site alone is sufficient to block generation of the pro-apoptotic p40MET fragment and phenocopies the apoptotic resistance observed in METex14Del-expressing cells. In contrast, deletion of the CBL-binding motif has no effect on apoptosis. However, deletion of both motifs simultaneously fully recapitulates METex14Del-driven tumorigenesis in HGF-humanized mice. Notably, despite driving as many tumors as METex14Del, the double mutation does not induce a rich or diverse transcriptional program, indicating that these tumors are independent of HGF-mediated signaling. Importantly, re-introduction of p40MET into METex14Del cells restores cell death sensitivity and robustly delay tumor growth, without altering MET signaling or migration. This demonstrates that loss of MET’s dependence receptor function is not merely correlative, but mechanistically required for tumor initiation. These findings are further supported by our NGS-based patient cohort analysis, in which we identified one tumor harboring a mutation affecting the caspase cleavage site, which may impair p40MET generation. However, we cannot exclude the possibility that this mutation might also interfere with phosphorylation of the regulatory tyrosine Y1003.

In 16HBE models, interestingly, loss of the CBL site appears to contribute to tumorigenesis through a mechanism at least partially independent of classical downstream signaling. Thus suggests the existence of additional, yet undefined, oncogenic responses. Notably, mutations affecting Y1003 have been reported in a few cancer patients who respond to MET-targeting TKIs. This supports the relevance of this motif in oncogenic MET activation^40–42^. Furthermore, in mammary epithelial cell lines overexpressing MET Y1003F, we have found this mutation to induce downstream signaling activation and cell motility in a manner comparable to METex14Del. This highlights the functional role of CBL binding site loss^13^.

Altogether, our study establishes MET as a unique example of a dual-function RTK whose tumorigenic potential is contingent on concomitant activation of oncogenic signaling and inactivation of a pro-apoptotic dependent receptor function. This challenges the longstanding paradigm of RTKs as purely oncogenic and introduces the concept that loss of tumor-suppressor activity may be an underappreciated contributor to cancer initiation progression.

This model echoes observations made for other dependent receptors such as TrkC, DCC, and UNC5H, where tumor-associated alterations —whether epigenetic or linked to ligand overexpression— bypass DR-mediated apoptosis to promote tumorigenesis^26,43^. In the case of METex14Del, this bypass occurs genetically, through deletion of the caspase cleavage site. This provides the first demonstration of a genetically encoded loss-of-function affecting a dependent receptor and contributing directly to human cancer. These findings open new avenues for targeting RTKs not only as oncogenes to be inhibited, but also as potential tumor suppressors to be reactivated or replaced by addition of an apoptotic therapy.

## EXPERIMENTAL MODEL AND STUDY PARTICIPANT DETAILS

### Patient cohorts

A total of 6,990 patient tumor samples from Lille University Hospital were analyzed with the CLAP next-generation sequencing (NGS) panel. Compliance with personal data protection regulations was ensured through a formal declaration submitted to the hospital’s Data Protection Officer (reference no. DEC25-244). In accordance with the General Data Protection Regulation (GDPR), patients were informed via routine medical correspondence —including information on potential data reuse— and/or through a dedicated information letter related to the study. Potential objections were verified by reviewing individual medical records and the Hospital Information System. A Privacy Impact Assessment was conducted prior to any data processing. All data were fully anonymized before collection, storage, and analysis. The methodology for NSCLC sample analysis adhered to the ethical principles outlined in the Declaration of Helsinki.

### Mouse models

All animal experiments were conducted in accordance with institutional and national ethical guidelines and were approved by the French Committee on Animal Experimentation and the French Ministry of Education and Research (authorization #50148-2024051317336526 v3). For xenograft studies with 16HBE cells, NOD.scid.Il2Rγc (NSG) mice carrying a humanized knock-in allele for HGF (originally developed by The Jackson Laboratory and bred by Charles River Laboratories) were used. For the xenograft study with Hs746T cells, SCID mice were purchased from the Institut Pasteur de Lille. Both male and female mice, aged 5 to 8 weeks, were included in the studies. Mice were housed at the PLETHA (Institut Pasteur de Lille) facility under specific pathogen-free conditions, with a 12-hour light/dark cycle and controlled ambient temperature maintained at 22 °C. All *in vivo* experiments with 16HBE cells were performed in an isolator with six mice housed in a M-BTM cage (Innovive) and allowed to eat and drink *ad libitum*.

### Cell lines

The 16HBE14o⁻ cell line was kindly provided by Prof. Dieter C. Gruenert (University of California) in 2016. A549 cells were a generous gift from Prof. Stéphanie Kermorgant (Barts Cancer Institute, London), and genome editing of these cells was performed as previously described (Hoggard et al., submitted). Hs746T cells were purchased from ATCC (Manassas, VA) in 2016. 16HBE14o⁻ parental cells and derived cell lines were cultured in Minimum Essential Medium (MEM; Invitrogen, Carlsbad, CA) supplemented with 10% heat-inactivated fetal bovine serum (FBS; Sigma-Aldrich, St. Louis, MO) and 1% ZellShield antibiotic mix (Minerva Biolabs, Berlin, Germany). A549 and Hs746T cells were maintained in Dulbecco’s Modified Eagle Medium (DMEM) GlutaMAX (Invitrogen), supplemented with 10% heat-inactivated FBS and 1% ZellShield. All cell lines were cultured at 37 °C in a humidified atmosphere containing 5% CO₂. Early-passage cell lines were routinely tested for mycoplasma contamination (MycoAlert, Lonza #LT07-318, Walkersville, MD), and all experiments were performed with cells maintained in culture for less than 3 months post-thawing. Cell line authentication was performed by short tandem repeat (STR) profiling (Eurofins Genomics, Germany).

## METHOD DETAILS

### Generation of mutant 16HBE14o⁻ cell lines by CRISPR–Cas9-mediated homologous recombination

#### sgRNA design and cloning

Single guide RNA (sgRNA) sequences were designed with the CRISPR Design Tool (http://crispr.mit.edu) to minimize off-target effects. Complementary oligonucleotides (0.5 nmol each, synthesized by Integrated DNA Technologies, Coralville, IA) were annealed at 95°C for 5 minutes in annealing buffer, this being followed by slow cooling to 55°C. Annealed oligos were ligated into the BbsI-digested pSpCas9(BB)-2A-Puro plasmid (#62988 Addgene, Watertown, MA) with T4 DNA ligase (New England Biolabs, Ipswich, MA) at 37°C for 90 minutes. Ligation mixtures were used to transform TAM1 chemically competent *E. coli* (Active Motif, Carlsbad, CA) and screening was done for ampicillin-resistant colonies. Plasmid DNA was extracted with a standard miniprep kit (#740588,5 Macherey-Nagel, Düren, NRW, Germany) and verified by Sanger sequencing.

#### Cell line editing and ssODN design

Point-mutant 16HBE14o⁻ cell lines (MET Y1003F, MET V1001A Y1003F, MET V1001A) and isogenic controls were generated by CRISPR–Cas9-mediated homologous recombination with single-stranded oligodeoxynucleotide (ssODN) templates containing the desired point mutation flanked by silent substitutions. These synonymous mutations served as a genetic barcode to enhance allele-specific PCR specificity as described^44^. The ssODN sequences are listed in Table S1.

#### Nucleofection and clonal isolation

Cells were electroporated with the Amaxa Nucleofector II device (Lonza, Walkersville, MD) with program D-027 and with the Cell Line Nucleofector Kit L (Lonza). Each reaction included 2 µg CRISPR– Cas9 plasmid and 2 µL of the corresponding ssODN (50 µM). Following electroporation, cells were pooled and incubated for 24 h prior to selection with puromycin (1 µg/mL) for 48 h. Surviving cells were plated by limiting dilution in 96-well plates to allow clonal expansion. Genomic DNA was extracted with the NucleoSpin® Tissue kit (#740952.250 Macherey-Nagel). Clones harboring the desired edits were identified by allele-specific RT-qPCR as previously described^44^, and confirmed by next-generation sequencing (NGS).

### Next-generation sequencing

Targeted next-generation sequencing was performed with a custom-designed panel partially based on the Ion AmpliSeq™ Colon and Lung Cancer Research Panel v2, with additional optimization to fully cover *MET* exon 14 and capture most known splice-site variants. The panel also included partial or complete exonic regions from key oncogenes and tumor suppressor genes frequently altered in solid tumors, including: *ACVR1, AKT1, ALK, BRAF, CTNNB1, EGFR, ERBB2, ERBB4, FGFR1, FGFR2, FGFR3, GNA11, GNAQ, GNAS, H3F3A, H3F3B, HIST1H3B, HIST1H3C, HRAS, IDH1, IDH2, KRAS, KIT, MAP2K1, MET, NRAS, PDGFRA, PIK3CA, PTEN, PTPN11, SMAD4*, and *TP53*. Library preparation was performed as previously described^45^.

All edited 16HBE cell lines displayed the expected editing on one allele, while the second allele carried a frameshift deletion leading to allele inactivation. Genome editing and MET exon 14 skipping in the two 16HBE clones were validated, as previously described^37^. Two independent METex14Del clones were previously established (cl.1 and cl.2), corresponding respectively to the published clone F and clone 7^37^.

### Generation of cell lines inducibly expressing p40MET

To generate stable inducible cell lines, the human p40MET coding sequence was cloned into the SfiI sites of the pSBtet-GH plasmid. This sleeping beauty (SB)-based vector enables doxycycline-inducible expression of p40MET, under the control of the synthetic RPBSA promoter^23^, alongside constitutive expression of the rtTA protein, a hygromycin resistance protein and a green fluorescent protein (GFP). Cloning of p40MET was performed with the In-Fusion cloning kit (Takara Bio Europe, Saint-Germain-en-Laye, France) according to the manufacturer’s protocol. Edited METex14Del 16HBE cells were co-transfected with the p40MET-pSBtet-GH construct and a sleeping beauty transposase expression vector (SB100X), a generous gift of Prof. Rolf Marschalek (Goethe-University of Frankfurt, Germany). The pSBtet-GH plasmid was a gift from Eric Kowarz^46^ (#60498 Addgene; http://n2t.net/addgene:60498; RRID: Addgene_60498). Stable cells were selected by culturing in hygromycin-containing medium for 10 days. Subsequently, highly GFP-positive cells were isolated by flow cytometry so as to enrich for stable p40MET production. The generated inducible cell line was then validated for p40MET expression in response to doxycycline treatment.

### Flow cytometry

Adherent cells were detached with StemPro Accutase (Life Technologies), washed with PBS, and incubated for 30 minutes at 4 °C with a primary antibody targeting the extracellular domain of MET (#AF276, R&D Systems; 2.5 µg per 10⁶ cells). After two PBS washes, cells were incubated for 20 minutes at 4 °C with an Alexa Fluor 647-conjugated secondary antibody (1:1000 dilution; #A-21469, Invitrogen), then resuspended in PBS at a concentration of 2 × 10⁶ cells/mL for flow cytometry analysis. Control samples were either unlabeled or incubated with secondary antibody only. Data acquisition was performed with an LSR Fortessa X20 flow cytometer (BD Biosciences), and analyses were carried out with BD FACSDiva™ and FlowJo software.

### Immunoblotting

For signaling studies, cells were treated with either DMSO (vehicle control), HGF at 30 ng/mL (#130-103-437 Miltenyi Biotec, Auburn, CA), or capmatinib at 1 µM (#HY-13404 MedChemExpress, Monmouth Junction, NJ). Cell pellets were lysed in PY lysis buffer (20 mM Tris-HCl pH 7.4, 50 mM NaCl, 5 mM EDTA, 1% Triton X-100, 0.02% sodium azide) supplemented with protease and phosphatase inhibitors: 1% aprotinin (Sigma-Aldrich, St. Louis, MO), 1 mM phenylmethylsulfonyl fluoride (PMSF; Sigma-Aldrich), 1 μM leupeptin (Sigma-Aldrich), 1 mM sodium orthovanadate (Na₃VO₄; Sigma-Aldrich), and 20 mM β-glycerophosphate (Sigma-Aldrich). Soluble protein extracts were quantified with the BCA Protein Assay Kit (#23227 Thermo Fisher Scientific, Waltham, MA). For immunoblotting, 25 μg total protein per sample was mixed with 3× NuPAGE LDS Sample Buffer (Invitrogen) to a final concentration of 1×, heated at 70 °C for 10 min, and resolved by SDS-PAGE on NuPAGE 4–12% Bis-Tris gels (Invitrogen). Proteins were transferred onto polyvinylidene difluoride (PVDF) membranes (Millipore, Burlington, MA), which were subsequently blocked for 1 h at room temperature in blocking buffer (0.2% casein and 0.1% Tween-20 in PBS). Membranes were incubated overnight at 4 °C with the indicated primary antibodies, washed three times in PBS containing 0.05% Tween-20, and incubated for 1 h at room temperature with species-specific horseradish peroxidase (HRP)-conjugated secondary antibodies. Following three additional washes in PBS/Tween, signal detection was performed with the SuperSignal West Dura chemiluminescent substrate (#34075 Thermo Fisher Scientific) and membranes were imaged with a ChemiDoc Imaging System (Bio-Rad, Hercules, CA). The primary antibodies and dilutions used were: mouse anti-MET (Cell Signaling Technology, Danvers, MA, #3148; 1:1000), rabbit anti-phospho-MET (Tyr1234/1235) (Cell Signaling Technology #3126; 1:1000), rabbit anti-phospho-MET (Tyr1003) (clone 13D11; Cell Signaling Technology #3135; 1:1000), rabbit anti-MET (clone SP44; Abcam, Cambridge, MA, #ab227637; 1:1000), rabbit anti-c-CBL (Cell Signaling Technology #2747; 1:1000), mouse anti-AKT (Cell Signaling Technology, #2920; 1:2000), rabbit anti-phospho-AKT (Ser473) (Cell Signaling Technology #4060; 1:2000), mouse anti-ERK2 (clone D2; Santa Cruz Biotechnology, Dallas, TX, #sc-1647; 1:1000), mouse anti-phospho-ERK1/2 (Thr202/Tyr204) (clone E10; Cell Signaling Technology, #9106; 1:1000), rabbit anti-PARP1 (clone H250; Santa Cruz Biotechnology #sc-7150; 1:1000), rabbit anti-cleaved caspase-3 (Asp175) (Cell Signaling Technology #9661; 1:1000), rabbit anti-Bak (Cell Signaling Technology #3814, 1:500), rabbit anti-Bax (Cell Signaling Technology #2774, 1:500), rabbit anti-Bcl-xL (clone 54H6, Cell Signaling Technology #2764, 1:1000), rabbit anti-Bid (Cell Signaling Technology #2002, 1:1000), rabbit anti-Bim (Cell Signaling Technology #2819, 1:500), rabbit anti-Mcl-1 (clone D35A5, Cell Signaling Technology #5453, 1:1000), mouse anti-Noxa (Calbiochem, San Diego, CA, #114C307, 1:250), rabbit anti-Puma (Cell Signaling Technology #4976, 1:500), mouse anti-actine (clone C4, Millipore #MAB1501, 1:4000), mouse anti-α-tubulin (clone B-5-1-2; Santa Cruz Biotechnology #sc-23948; 1:1000), mouse anti-GAPDH (clone 6C5; Santa Cruz Biotechnology #sc-32233). Species-specific horseradish peroxidase (HRP)-conjugated secondary antibodies were used at a 1:10000 dilution: donkey anti-rabbit IgG (Jackson ImmunoResearch, West Grove, PA, #711-035-152) and goat anti-mouse IgG (Jackson ImmunoResearch #115-035-146). Quantification of immunoblot band intensities was performed with LabFigures.com (Sciugo.com) or ImageQuant software.

### Wound healing assay and invasion assays

Cells (50 000/mL) were seeded into 96-well ImageLock plates (Sartorius, Göttingen, Germany) and grown to full confluence. The following day, cells were serum-starved for 2 h. For migration assays, serum starvation was done in the presence of mitomycin C (1 μg/mL; Sigma-Aldrich) to inhibit proliferation. A uniform scratch was generated with the WoundMaker tool (Essen BioScience, West Wickham, Kent, UK), and wells were rinsed twice with medium to remove cellular debris. Wound closure was monitored every 2 h for up to 8 h for migration or 72 h for invasion with the Incucyte SX5 Live-Cell Analysis System (Sartorius). Cell migration was quantified with the Incucyte software based on relative cell density within the wound area, normalized to the initial timepoint (t₀).

For invasion assays, cells were seeded onto a layer of Growth Factor Reduced Matrigel (Corning, Corning, NY) diluted 1:100 in PBS. After scratch formation, an additional layer of Matrigel diluted 1:1 (v/v) in medium was applied. Imaging and quantification were performed as described for the migration assay.

### Scattering assays

Cells were seeded at low density (∼1,500 cells per well) into 24-well plates and allowed to adhere overnight. After 48 h, the plates were transferred to the Incucyte SX5 Live-Cell Analysis System (Sartorius), and phase-contrast images were acquired every 6 h for 96 h with the Cell-by-Cell Adherent Cell Module. Segmentation images were exported and analyzed with a custom ImageJ macro based on particle analysis, with a size filter applied to exclude debris and non-cellular objects. For each timepoint, individual cell clusters were cropped, and the x and y coordinates of each cell were extracted on the basis of the centroid position. Cell density was computed with a custom R script. Dispersion was quantified as the inverse of local cell density at defined timepoints.

### Cell viability assay

Cells were seeded at a density of 2,500 cells per well into 96-well plates in medium supplemented with 0.5% FBS. Real-time monitoring of cell growth and confluence was performed with the Incucyte SX5 Live-Cell Analysis System (Sartorius). At the experimental endpoint, cell viability was assessed with the Alamar Blue Cell Viability Reagent (Invitrogen), following an 8 h incubation at 37°C. Absorbance was measured at 570 nm and 590 nm with a MultiSkan RC plate reader (Thermo Labsystems, Helsinki, Finland) and metabolic activity was quantified as a proxy for cell viability.

### Apoptosis assay

Cells were seeded at a density of 2,500 cells per well into 96-well plates in medium containing 0.5% FBS. Apoptotic cell death was assessed in real time with the Incucyte SX5 Live-Cell Analysis System (Sartorius). Three complementary fluorescent readouts were used: Annexin V staining (#4641 Annexin V Green Reagent for Apoptosis, Sartorius) to detect phosphatidylserine externalization, caspase 3/7 activity (#4440 Caspase 3/7 Green Apoptosis Assay Reagent, Sartorius), and membrane permeability (#4632 Cytotox Red Reagent, Sartorius). Fluorescence signal confluence was quantified and normalized to the total confluence in each well to take variations in cell density into account. During serum deprivation experiments, images were acquired every 4 hours. In selected experiments, cells were treated with 1 µM staurosporine (#282T6680 Tebubio, Le Perray-en-Yvelines, France) for 24 h to induce acute apoptosis, with image acquisition every hour.

### Immunofluorescence staining

Cells were seeded onto glass coverslips in 12-well plates at a density of 100,000 cells per well. After 48 h of culture, the cells were serum-starved for 2 h and subsequently subjected to the indicated treatments. Following treatment, the cells were washed with PBS and fixed in cold methanol:acetone (1:1, v/v). Permeabilization was carried out with PBS containing 0.5% Triton X-100, followed by a 30-minute blocking step in PBS supplemented with 0.2% casein (w/v). Coverslips were incubated with Rabbit Anti-Cleaved Caspase 3 (Asp175) primary antibody (Cell Signaling Technology #9661; 1:400) for 1 h at room temperature, washed with PBS, and then incubated for 1 h with Alexa Fluor 488-conjugated secondary antibodies (#A11034 Anti-Rabbit IgG, 1:1000; Sigma-Aldrich). Nuclei were counterstained with Hoechst 33258 (Sigma-Aldrich). Samples were imaged with a Zeiss AxioImager Z1 microscope equipped with an ApoTome module (Plan-Apochromat objective, NA 1.40, oil immersion) and ZEN acquisition software (Zeiss).

### *In vitro* caspase 3 cleavage assay

Cells were lysed in a caspase-activity-compatible buffer (20 mM PIPES, pH 7.2, 100 mM NaCl, 1% CHAPS, 10% sucrose, 5 mM DTT, 50 µM EDTA), and lysates were clarified by centrifugation. Cleavage reactions were performed by incubating the lysates with 1 µL purified recombinant active caspase 3 (kind gift from the Burnham Institute) for 4 h at 37 °C. Reactions were terminated by addition of SDS sample buffer, boiled, and analyzed by immunoblotting to detect cleavage of target proteins.

### Transcriptomic analysis

Cells were seeded into complete medium (750,000 cells per dish) and serum-starved the following day for 2 hours before stimulation with or without 30 ng/mL HGF in serum-free medium for 24 hours. Total RNA was extracted with the Nucleospin RNA kit (Macherey-Nagel) according to the manufacturer’s instructions, including on-column DNase I treatment to remove genomic DNA. HGF stimulation and RNA extraction were performed in four independent experiments. RNA concentration and purity were assessed with the NanoDrop 2000c spectrophotometer (ThermoScientific). Construction of sequencing libraries using the Quants-Pool Sample-Barcoded 3’mRNA-Seq Library Prep for Illumina (Lexogen): Briefly, starting with 6 µL containing a total of 40 ng of RNA, we add 1 µL of ERCC spike-in control. Library preparation begins with oligo(dT) priming, where the primer includes both a sample-specific index for sample multiplexing and a Unique Molecular Identifier (UMI). Samples are then pooled in groups of 24, including one control sample per pool. Following purification and RNA digestion, second-strand synthesis is initiated using random priming. Once the double-stranded cDNA library is generated, it is purified using magnetic beads and amplified through 11 PCR cycles. The final library **is** purified and quality-checked using the Agilent Bioanalyzer 2100. Each library is then pooled equimolarly, and the final pool is also assessed on the Agilent Bioanalyzer 2100 before sequencing on the Illumina NovaSeq 6000 using 100-cycle chemistry, yielding between 8.2 million and 27 million reads per sample.

### Data analysis

The idemuxCPP (v0.3.0) tool was employed to demultiplex paired-end fastq files. To eliminate poor quality regions and poly(A) of the reads, we used the *fastp* program. We used a quality score threshold of 20 and removed reads shorter than 25 pb. Read alignments were performed with the *STAR* (v2.6.0a) program with the human reference genome (GRCh38) and the reference gene annotations (*Ensembl*). The UMI (Unique Molecular Index) allowed reducing errors and quantitative PCR bias with *fastp* (v0.24.0) and *umi-tools* (v0.5.4). On the basis of read alignments, we counted the number of molecules per gene with *FeatureCount* (v1.6.0). Other programs run for the quality control of reads and for the workflow included *qualimap* (v2.2.1), *FastQC* (v0.11.9), *FastQScreen* (v0.15.2), and *MultiQC* (v1.21). Differential gene expression analysis of RNA-seq data was performed with *R* (R version 4.2.2 (2022-10-31)), *Bioconductor* (v3.16) and package *DESeq2* (v1.38.3). The cut-off for differentially expressed genes was p-value padj (BH) < 0.05. *Over-Representation Analysis* (ORA) and *Gene Set Enrichment Analysis* were performed with the R library *clusterProfiler* (v4.6.0).

### Xenograft tumors

A total of 2 × 10⁶ cells suspended in PBS were injected subcutaneously into the left and right inguinal flanks of 5-to 8-week-old NSG-huHGF mice (Jackson Laboratory; bred by Charles River, Wilmington, MA), without distinction of sex. Tumor growth was monitored by measuring tumor length (L) and width (W) with digital calipers, and tumor volume was estimated with the standard formula: V = (L × W²)/2. For the capmatinib efficacy study, mice were randomized into treatment groups when the tumors reached an average size of ∼100 mm³. Capmatinib (#HY-13404 MedChemExpress) was freshly resuspended in vehicle (1% methylcellulose, 0.5% Tween-80 in water) at 2 mg/mL. The suspension was pH-adjusted to 2.5–3.5, sonicated (10 × 5 s) in an ultrasonic bath, and stored in the dark at 4°C. Mice received a daily oral gavage of either capmatinib (10 mg/kg/day for 16HBE cells or 2.5 mg/kg/day for Hs746T cells) or vehicle. Tumor dimensions and body weight were recorded biweekly to monitor progression and systemic toxicity. Mice were euthanized when one of the tumors reached 1000 mm³. All engrafted mice were included in the analysis; no data were excluded. Survival was assessed by the Kaplan–Meier method. Upon sacrifice, approximately 50% of each tumor was fixed in 4% paraformaldehyde for 24 h and then stored in 70% ethanol for histological analysis, while the remaining 50% were frozen at −80°C. For the inducible p40MET expression study, doxycycline (#9891 Sigma-Aldrich) was administered at 1 mg/mL in drinking water, refreshed every 48 hours throughout the course of the experiment.

### Immunohistochemistry

Tumors were fixed overnight in 4% paraformaldehyde (PFA), dehydrated through a graded ethanol series (30%, 70%, 95%, and 100%), cleared in toluene, and embedded in paraffin. Tissue sectioning was performed by the PLBS – BICeL core facility (University of Lille). Antigen retrieval was performed by heating until boiling for three times 1 min 30 s in Tris-EDTA pH 9.0 and the cooled down at room temperature for 30 min, followed by 1 h of protein blocking using a solution of BSA 5%, donkey sera 5%, Triton X100 0.3% to prevent non-specific binding. Immunostaining was carried out on deparaffinized sections with two primary antibodies: rabbit anti-MET (clone SP44; Abcam #ab227637; 1:1000) and mouse anti-PCNA (Abcam #ab19197; 1:1000), both diluted in blocking buffer, and incubated overnight at 4°C. Alexa Fluor 488-conjugated anti-rabbit IgG secondary antibody (Sigma-Aldrich #A11034; 1:2000) was applied for 1 h at room temperature. Nuclear counterstaining was performed with Hoechst 33258 (Sigma-Aldrich). Slides were imaged with a Zeiss AxioImager Z1 microscope equipped with an ApoTome module (Plan-Apochromat objective, NA 1.40, oil immersion) and ZEN acquisition software (Zeiss).

### Statistical analysis

Data are presented as means ± standard deviation (SD) or standard error of the mean (SEM), as indicated. Depending on the experimental design and data distribution, comparisons between groups were performed by one-way or two-way analysis of variance (ANOVA), followed by appropriate post hoc tests where applicable. All statistical tests were two-sided, with a significance threshold set at *p* < 0.05. Analyses were conducted with GraphPad Prism software, version 10 (GraphPad Software, San Diego, CA).

## AUTHOR CONTRIBUTIONS

R.T.: conceptualization, methodology, validation, formal analysis, investigation, data curation, writing – original draft, visualization. M.Fe.: conceptualization, methodology, validation, investigation, data curation, resources, writing – review & editing. A.V., S.P., A.L., and C.Vi.: methodology, investigation. E.W.: methodology, software. J.P.M.: formal analysis, visualization. C.Vu.: methodology, investigation, formal analysis. A.C.L: methodology. Z.K.: methodology, writing – review & editing. M.J.T.: methodology, investigation. C.D.: methodology, investigation, formal analysis, writing – review & editing. L.G.: methodology, resources, writing – review & editing. A.C., E.W.: methodology, investigation. M.Fi.: formal analysis, methodology, resources, software, supervision, visualization. S.K.: methodology, resources. L.P.: conceptualization, funding acquisition, methodology, resources, supervision, writing – review & editing. A.P.: conceptualization, investigation, methodology, resources. D.T.: conceptualization, funding acquisition, methodology, project administration, supervision, validation, visualization, writing – review & editing.

## Acknowledgments

We thank the US 41 – UAR 2014 - PLBS platform (University of Lille) for support with flow cytometry and histological processing, including tissue embedding and sectioning, and the US 41 – UAR 2014 - PLETHA animal facility at Institut Pasteur de Lille for mouse housing and care.

This article contains supporting information.

## RESOURCES AVAILABILITY

### Lead contact

Further information and requests for resources, reagents, and data should be directed to the Lead Contact, David Tulasne, who will fulfill reasonable requests.

### Materials availability

Materials and reagents are available from Lead Contact upon reasonable request.

---

**Figure S1.**
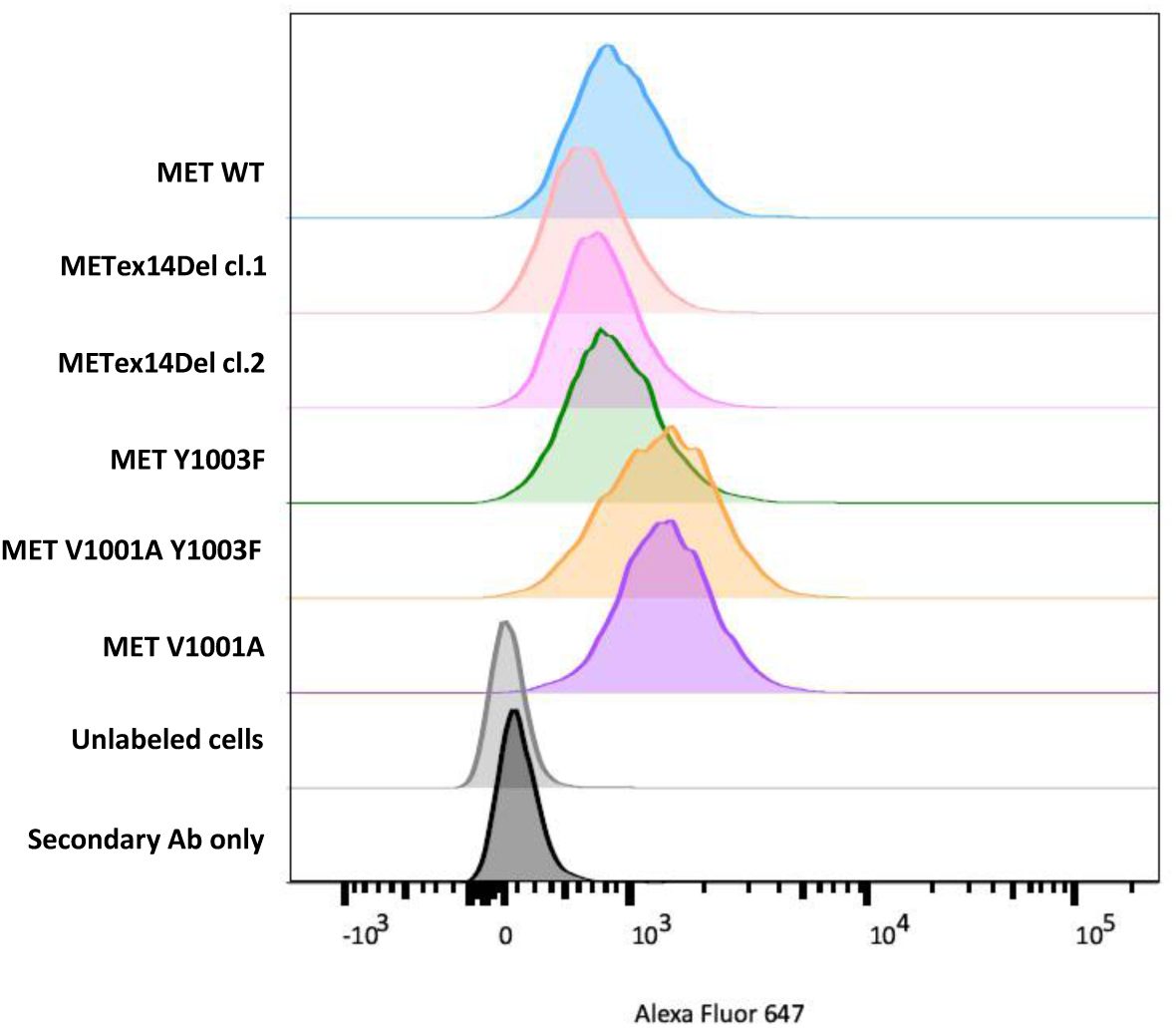
MET-mutated-16HBE cell line validation by flow cytometry. Plasma-membrane MET expression of was evaluated by flow cytometry with an antibody targeting the extracellular domain of the receptor.

**Figure S2.**
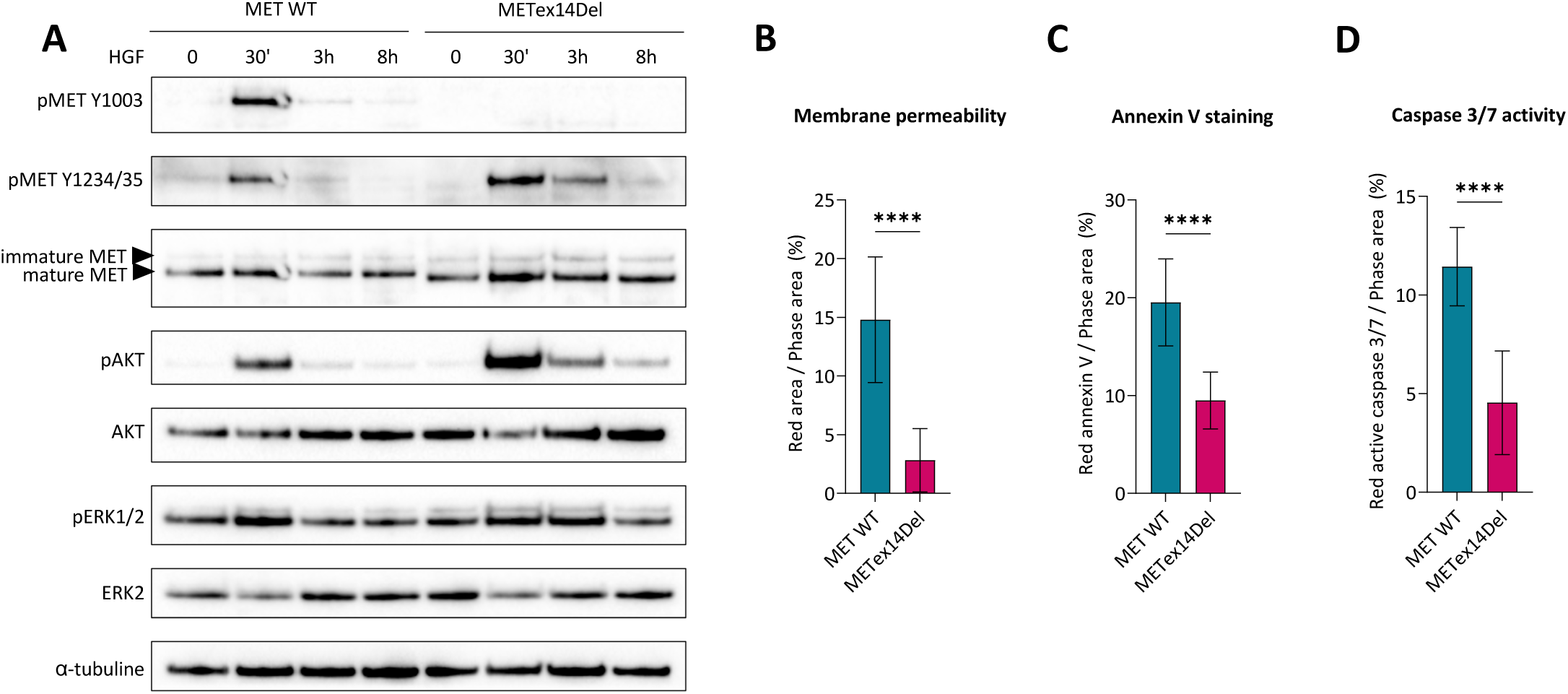
METex14Del induces sustained signaling and resistance to apoptosis in A549 cells. (A) Immunoblot analysis of MET-induced downstream signalling pathways in edited A549 cells after treatment with 30 ng/mL HGF for the indicated time. Immunoblotting is representative of three biological replicates. (B-D) Membrane permeability (B), Annexin V staining (C), and active caspase 3/7 staining (D) were measured after seven days of serum starvation. Fluorescence signals were normalized to cell confluence. *n=5;* mean ± SD; representative of three independent experiments. Statistical significance was determined by one-way ANOVA. **** (*p* < 0.0001).

**Figure S3.**
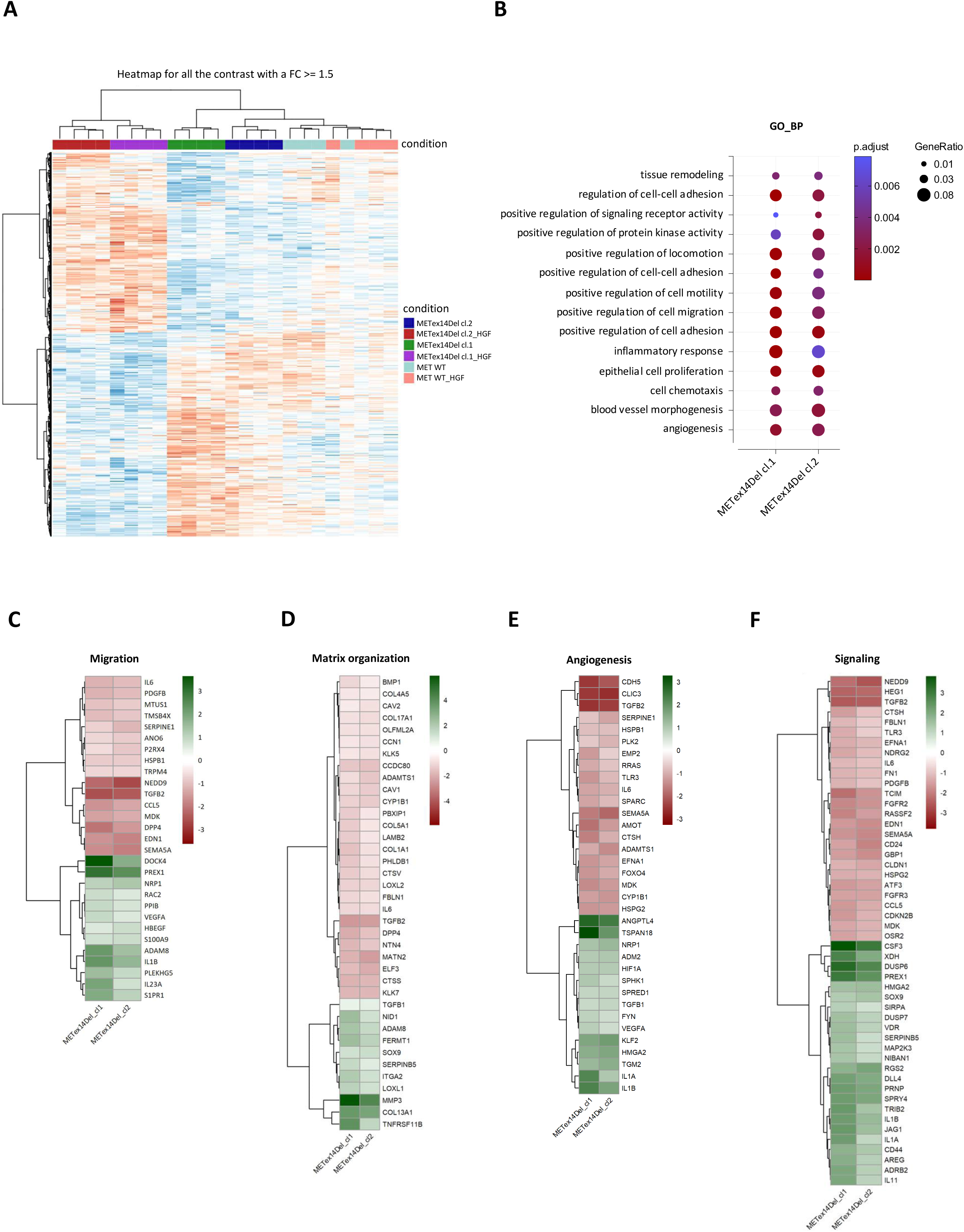
Transcriptomic profile of METex14Del 16HBE clones compared to MET WT. (A) Heatmap showing differentially expressed genes across all pairwise contrasts (absolute fold change ≥ 1.5; adjusted *p*-value (BH) < 0.05). (B) Gene ontology (GO) Biological Process (BP) enrichment analysis of differentially expressed genes between each METex14Del clone and its control represented as a dot plot. Selected GO terms of interest are highlighted; dot size indicates gene ratio and color represents adjusted *p*-value. Differential expression and GO analysis were performed with the DESeq2 and clusterProfiler package. Threshold used: |fold change| ≥ 1.5 and adjusted *p*-value (BH) < 0.05. (C–F) Clustered expression profiles of genes modulated in both METex14Del clones in response to HGF within selected GO categories: cell migration (C), extracellular matrix organization (D), angiogenesis (E), and cell signaling (F). Differential expression and GO analyses were performed with the DESeq2 and clusterProfiler package. Thresholds used: |fold change| ≥ 1.5 and adjusted *p*-value (BH) < 0.05.

**Figure S4.**
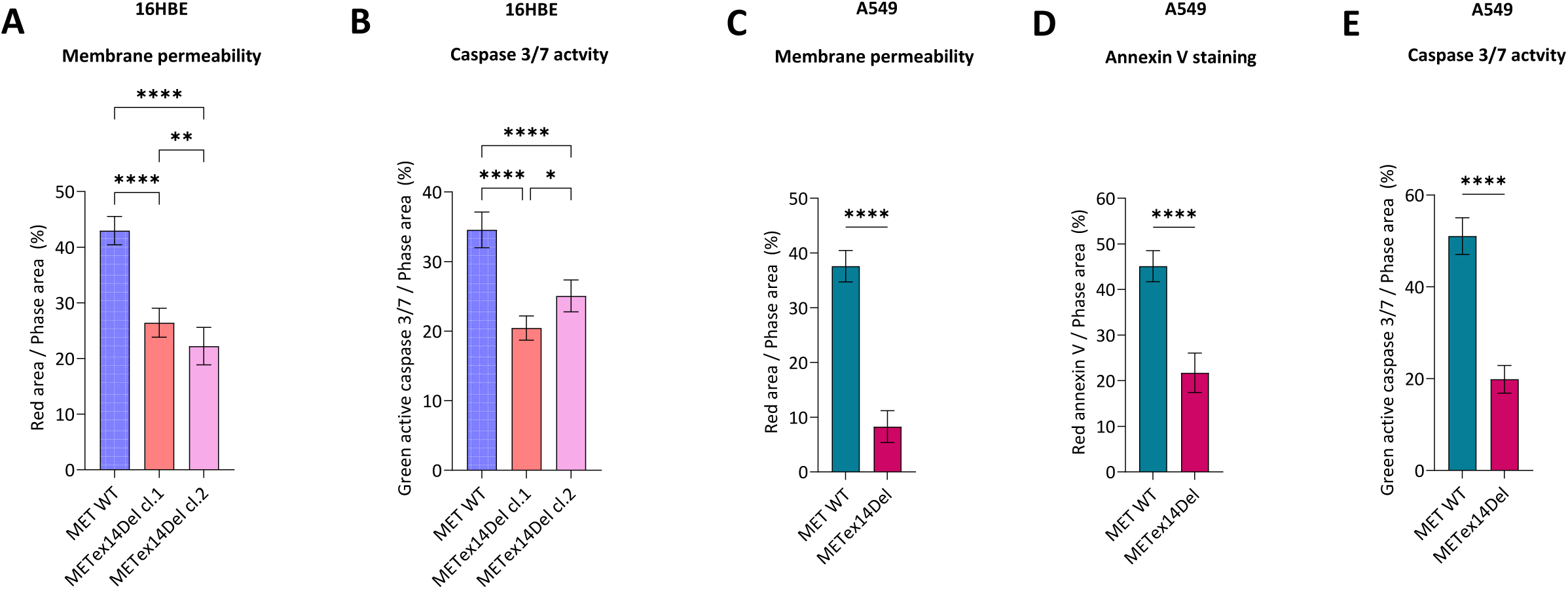
METex14Del induces resistance to short-term apoptosis in both 16HBE and A549 cells. (A-B) Membrane permeability (A), and active caspase 3/7 staining (B) were measured in edited 16HBE cells after 8 h of staurosporine treatment at 1 µM. Fluorescence signals were normalized to cell confluence. *n=5;* mean ± SD; representative of three independent experiments. Statistical significance was determined by one-way ANOVA. * (*p* < 0.05), ** (*p* < 0.01), **** (*p* < 0.0001). (C-E) Membrane permeability (C), Annexin V staining (D), and active caspase 3/7 staining (E) were measured 24 h after staurosporine treatment at 1 µM. Fluorescence signals were normalized to cell confluence. *n=5;* mean ± SD; representative of three independent experiments. Statistical significance was determined by one-way ANOVA. **** (*p* < 0.0001).

**Figure S5.**
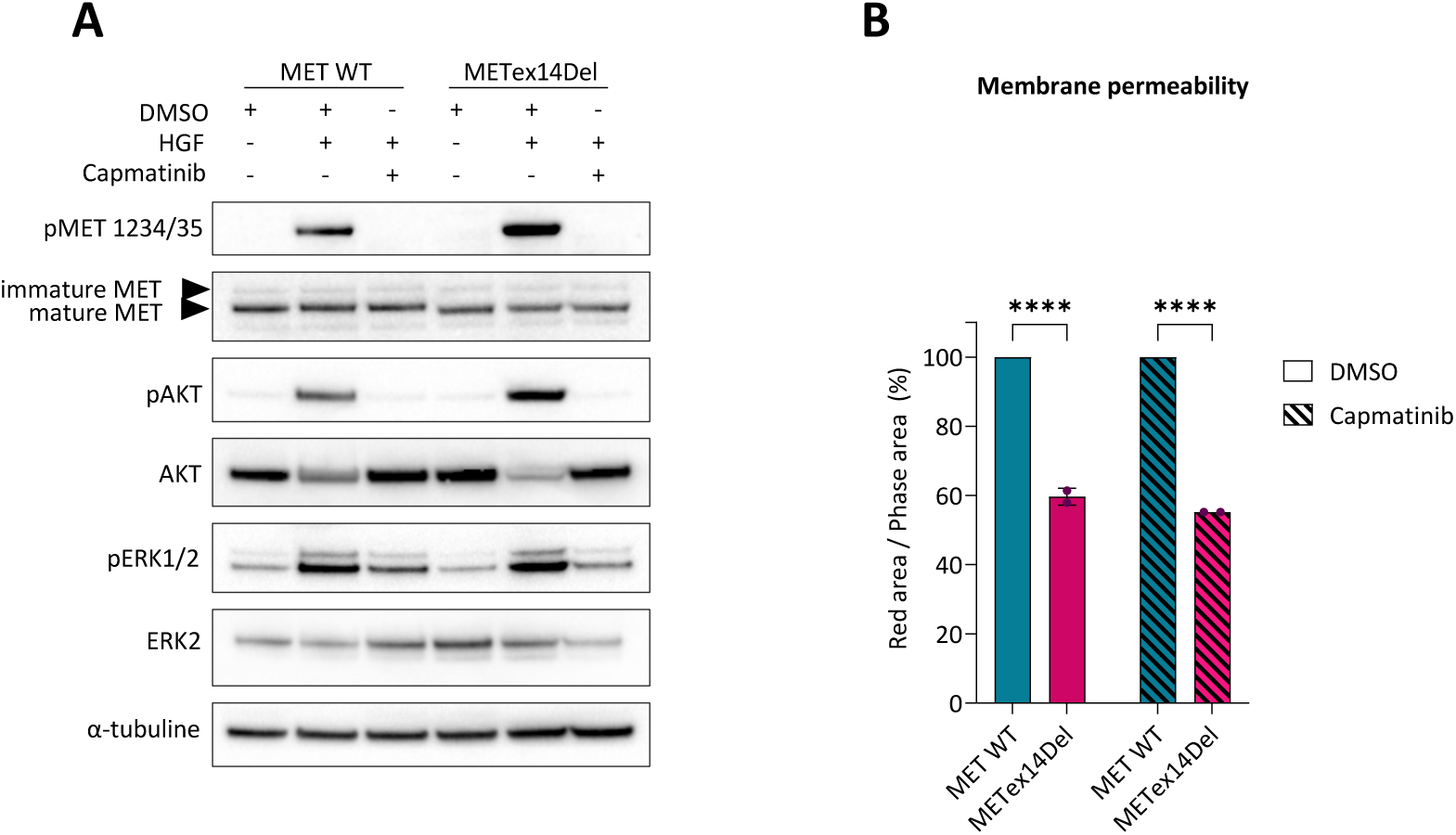
Capmatinib inhibits METex14Del-induced signaling in A549 cells. (A) Immunoblot analysis of MET-induced downstream signalling pathways in A549 editing cells treated with 30 ng/mL HGF for 30 minutes, with or without capmatinib pre-treatment. Cells were pre-treated with 1 µM capmatinib for 90 minutes prior to HGF stimulation. Immunoblots are representative of three independent biological experiments. (B) Membrane permeability was measured five days after serum starvation with or without 1 µM of capmatinib. Fluorescence signal was normalized to cell confluence. Data represent means ± SD from two independent biological experiments, each including at least six technical replicates. Statistical significance was determined by one-way ANOVA. **** (*p* < 0.0001).

**Figure S6.**
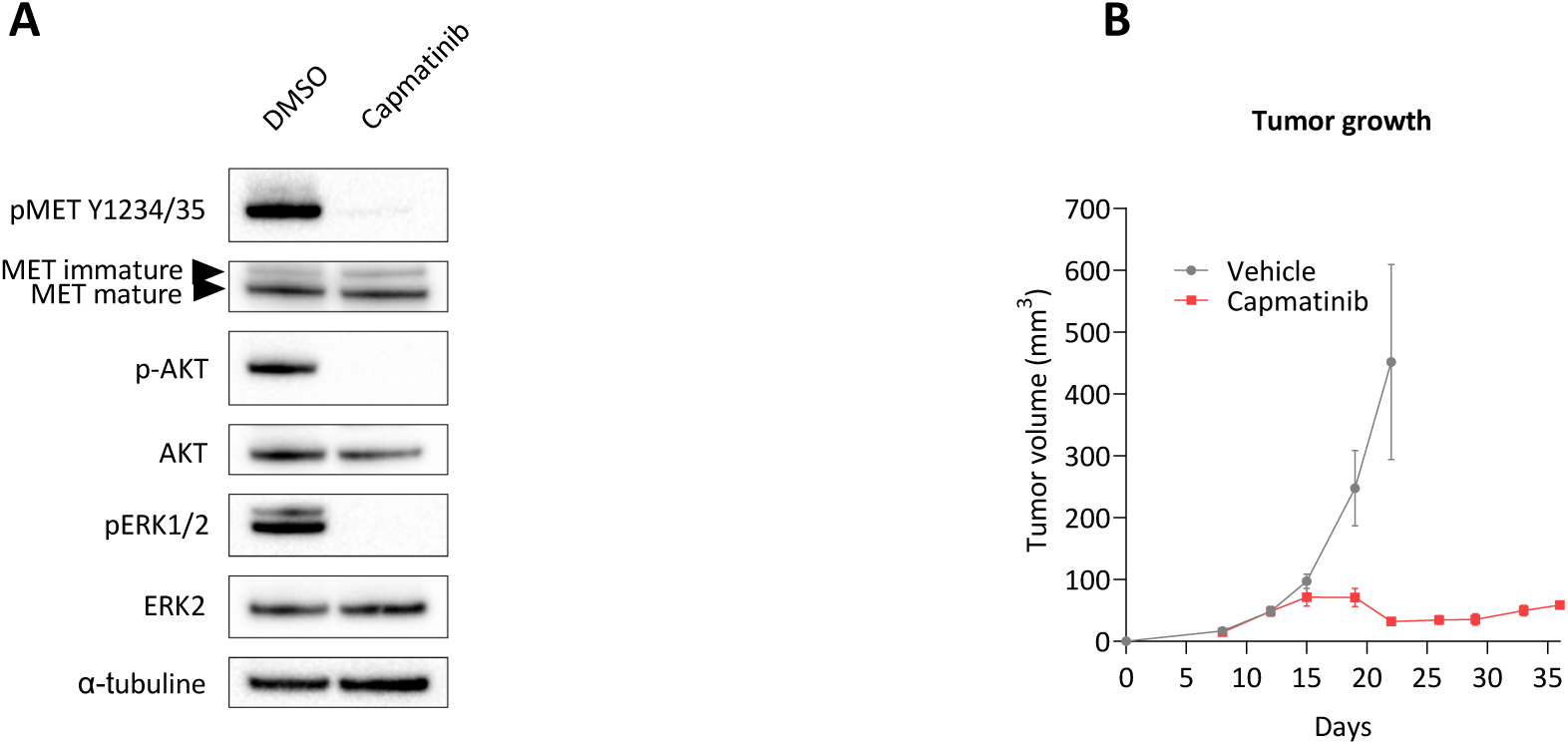
Capmatinib inhibits METex14Del-induced tumorigenesis in Hs746T cells. (A) Tumor growth of SCID mice injected subcutaneously with Hs746T cells. Daily treatment with vehicle or capmatinib (2.5 mg/kg/day) was initiated fifteen days after cells injection. Data represent mean tumor volume ± SEM. (B) Immunoblot analysis of MET-induced downstream signalling pathways in Hs746T treated with or without capmatinib 1 µM for 2 h. Immunoblots are representative of two independent biological experiments.

**Figure S7.**
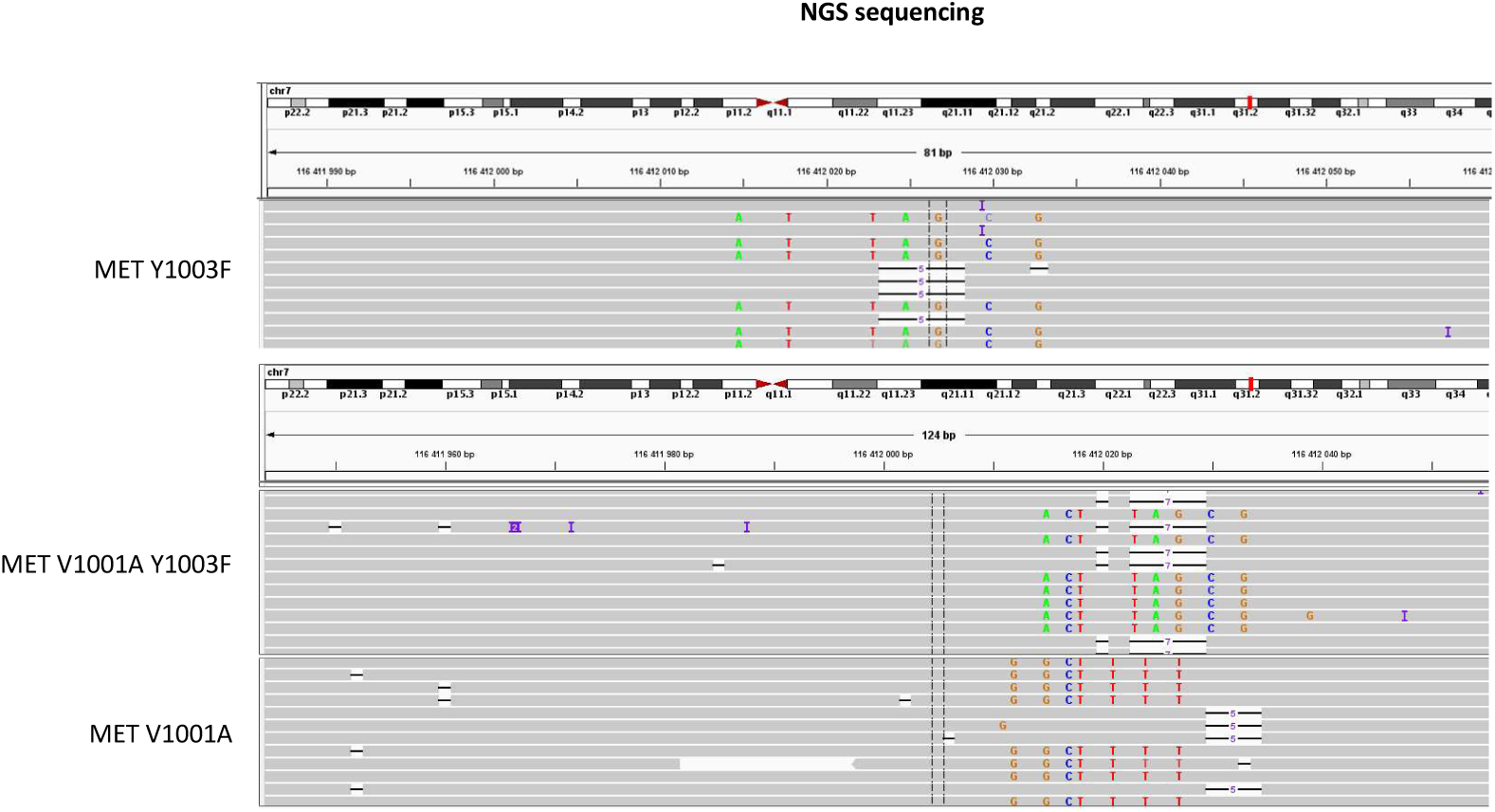
MET mutated-16HBE cell lines validation by high-throughput sequencing. Result of targeted NGS of the edited 16HBE-cell genomes. For 16HBE MET Y1003F, a set of sequences shows insertion of the silent mutations and the mutation predicted to cause the Y1003F substitution (in color); another set of sequences shows a 5-base deletion leading to a frame shift. For 16HBE MET V1001A Y1003F, a set of sequences shows insertion of the silent mutations and the mutations predicted to cause the V1001A and Y1003F substitutions (in color); another set of sequences shows a 1-base and 7-base deletion leading to a frame shift. For 16HBE MET V1001A, a set of sequences shows insertion of the silent mutations and the mutations predicted to cause the V1001A substitutions (in color); another set of sequences shows a 5-base deletion leading to a frameshift.

**Figure S8.**
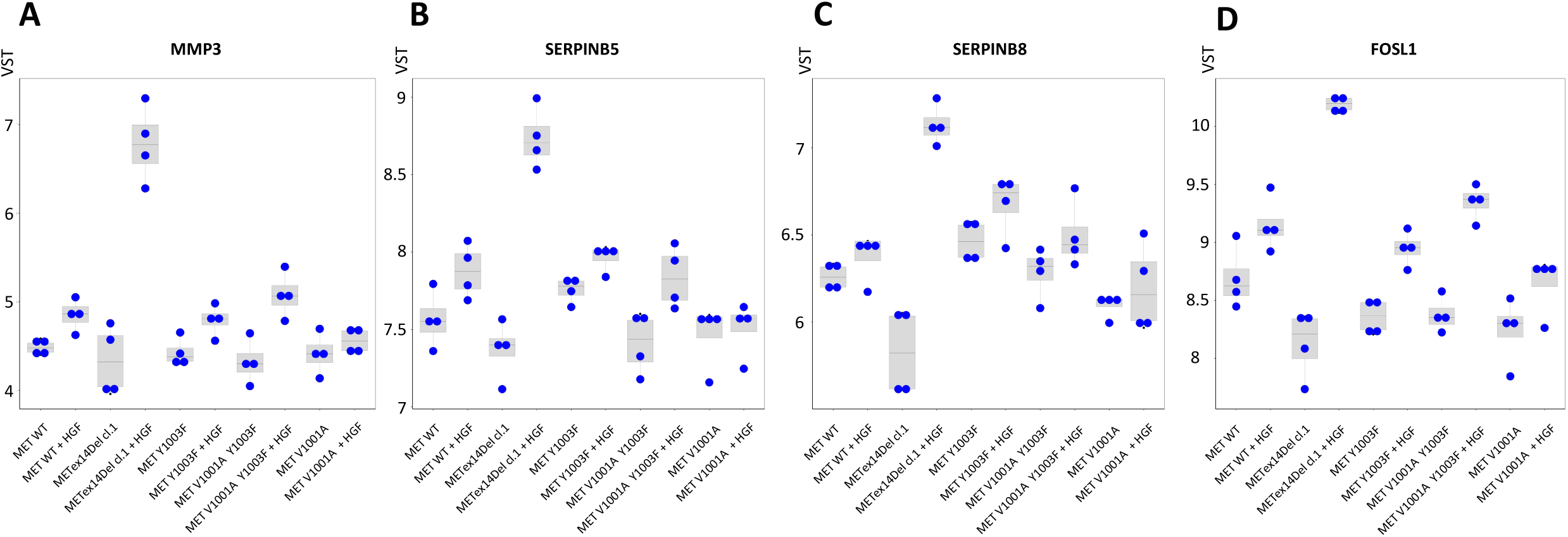
Expression of selected genes from the transcriptomic analysis. (A-D) Normalized expression of selected genes from RNA sequencing in genome-edited 16HBE clones. Variance-stabilized transformed (VST) counts are shown for MMP3 (A), SerpinB5 (B), SerpinB8 (C), and FOSL1 (D) across the different 16HBE edited clones, their control and +/- HGF. n=4 independent experiments

**Figure S9.**
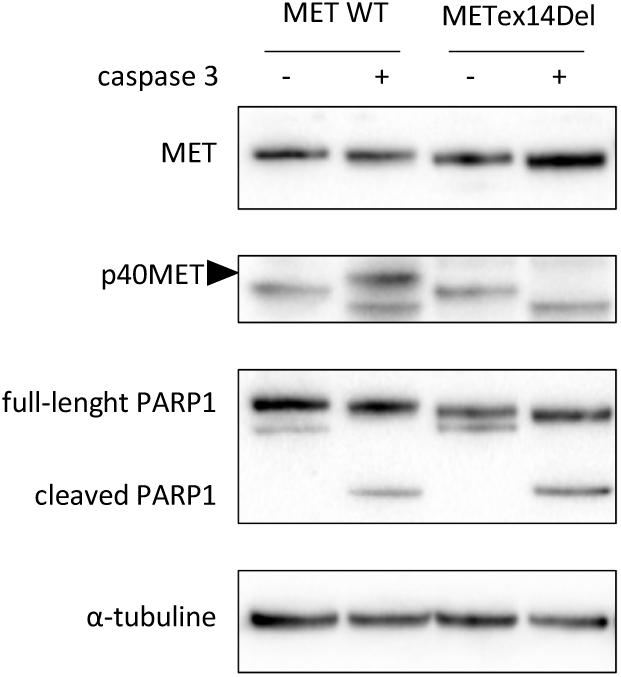
*In vitro* caspase 3 cleavage in A549 cells. Immunoblot analysis showing generation of the p40MET fragment following *in vitro* cleavage of MET by recombinant caspase 3 in A549 cell lysates. Immunoblots are representative of three independent biological experiments.

**Figure S10.**
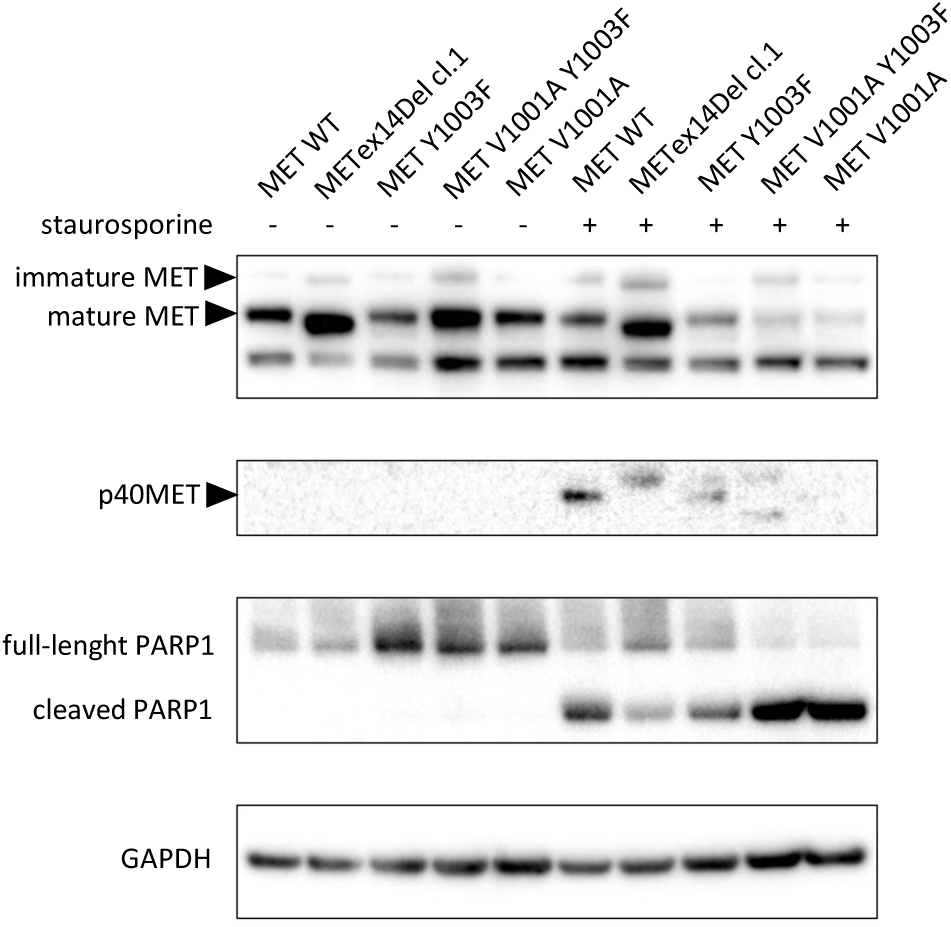
Staurosporine treatment induces p40MET fragment generation in 16HBE cells. Immunoblot analysis showing generation of the p40MET fragment in edited 16HBE cells after treatment with 5 µM staurosporine for five hours. Immunoblots are representative of three independent biological experiments.

**Figure S11.**
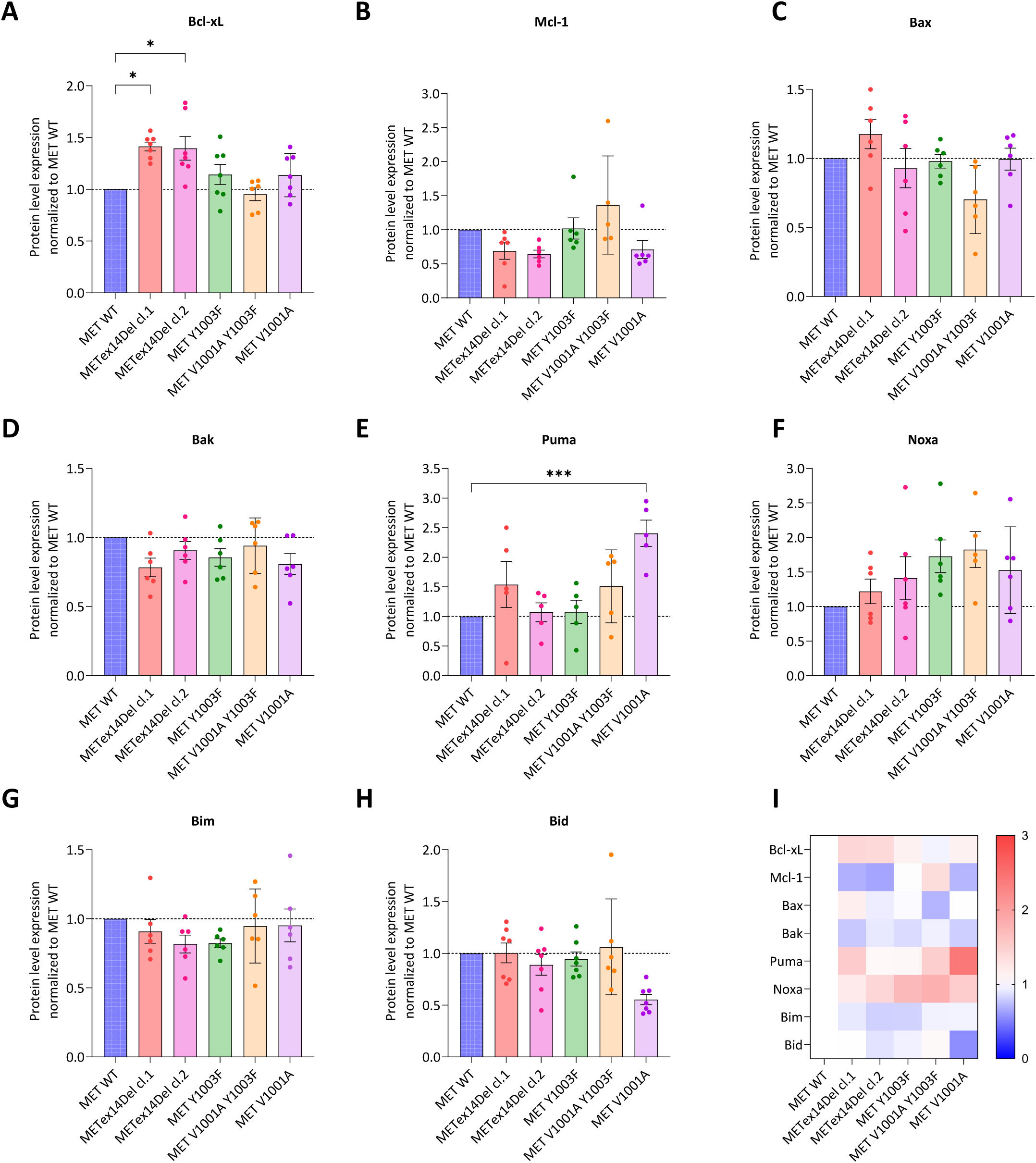
Expression levels of apoptosis-related proteins. Expression of apoptosis-related proteins in genome-edited 16HBE clones. (A–G) Quantification of Bcl-xL (A), Mcl-1 (B), Bax (C), Bak (D), Puma (E), Noxa (F), Bim (G), and Bid (H). Protein levels were normalized to β-actin and quantified from at least five independent blots. Data represent means ± SEM. (H) Heatmap summarizing the levels of all proteins across the clones. Statistical significance was determined by one-way ANOVA. ns (not significant), * (p < 0.05), ** (p < 0.01), *** (p < 0.001), **** (p < 0.0001).

**Supplementary Table S1:**
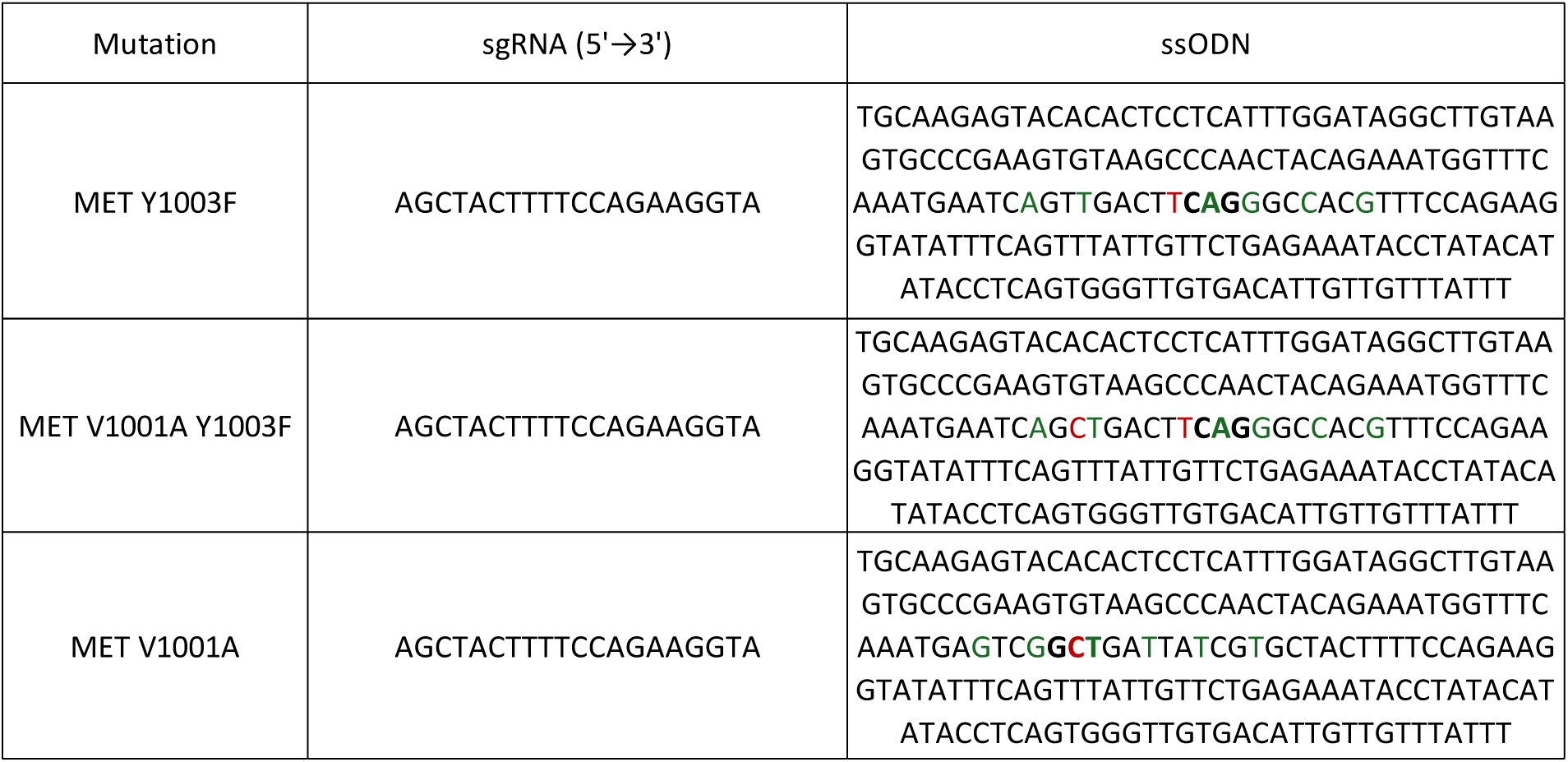
Sequence of ssODN for CRISPR-Cas9 editing. Silent mutations are represented in green and point mutations are represented in red.

